# Liprin-α/RIM complex regulates the dynamic assembly of presynaptic active zones via liquid-liquid phase separation

**DOI:** 10.1101/2024.08.29.610253

**Authors:** Gaowei Jin, Joaquín Campos, Yang Liu, Berta Marcó de la Cruz, Shujing Zhang, Mingfu Liang, Kaiyue Li, Xingqiao Xie, Cong Yu, Fredrik H. Sterky, Claudio Acuna, Zhiyi Wei

**Affiliations:** Shenzhen Key Laboratory of Biomolecular Assembling and Regulation, Southern University of Science and Technology, Shenzhen, China; Brain Research Center, Southern University of Science and Technology, Shenzhen, China; School of Life Sciences, Southern University of Science and Technology, Shenzhen, China; Chica and Heinz Schaller Foundation, Institute of Anatomy and Cell Biology, University of Heidelberg, Heidelberg, Germany; Department of Laboratory Medicine, Institute for Biomedicine, Sahlgrenska Academy, University of Gothenburg, Gothenburg, Sweden; Wallenberg Centre for Molecular and Translational Medicine, University of Gothenburg, Gothenburg, Sweden; Guangdong Provincial Key Laboratory of Cell Microenvironment, Southern University of Science and Technology, Shenzhen, China; Institute for Biological Electron Microscopy, Southern University of Science and Technology, Shenzhen, China; Department of Clinical Chemistry, Sahlgrenska University Hospital, Gothenburg, Sweden

## Abstract

Presynaptic scaffold proteins, including liprin-α, RIM, and ELKS, are pivotal to the assembly of the active zone and regulating the coupling of calcium signals and neurotransmitter release, yet the underlying mechanism remains poorly understood. Here, we determined the crystal structure of the liprin-α2/RIM1 complex, revealing a multifaceted intermolecular interaction that drives the liprin-α/RIM assembly. Neurodevelopmental disease-associated mutations block the formation of the complex. Disrupting this interaction in neurons impairs synaptic transmission and reduces the readily releasable pool of synaptic vesicles. Super-resolution imaging analysis supports a role for liprin-α in recruiting RIM1 to the active zone, presumably by promoting the liquid-liquid phase separation (LLPS) of RIM1. Strikingly, the liprin-α/RIM interaction modulates the competitive distribution of ELKS1 and voltage-gated calcium channels (VGCCs) in RIM1 condensates. Disrupting the liprin-α/RIM interaction significantly decreased VGCC accumulation in the condensed phase and rendered release more sensitive to the slow calcium buffer EGTA, suggesting an increased physical distance between VGCC and vesicular calcium sensors. Together, our findings provide a plausible mechanism of the liprin-α/RIM complex in regulating the coupling of calcium channels and primed synaptic vesicles via LLPS for efficient synaptic transmission and uncover the pathological implication of liprin-α mutations in neurodevelopmental disorders.

## Introduction

Synaptic transmission is the cornerstone of brain functions, representing the fundamental process through which neurons communicate. Triggered by an action potential, neurotransmitter release occurs at a specialized region in the presynaptic terminal, known as “active zone” ^1–3^. This dynamic region arises from the orchestrated assembly of a diverse array of proteins, forming an electron-dense structure attached to the plasma membrane that governs synaptic vesicle exocytosis. Five conserved scaffold proteins have emerged as core components in active zone assembly, including liprin-α, RIM, RIM-BP (RBP), ELKS, and Munc13 ^1^. However, the assembly mechanism remains poorly understood, mainly due to the limited understanding of the complex interactions among these proteins.

Among these core scaffolds, liprin-α has garnered increasing attention for its evolutionary conserved roles in active zone formation and function. The liprin-α family contains four members (liprin-α1/2/3/4) in mammals and one member each in *C. elegans* and *Drosophila* ^4–8^. Genetic studies in invertebrates reveal that dysfunction of liprin-α orthologs leads to altered active zone morphology and diminished synaptic vesicle accumulation ^5,6,9^. Although the depletion of two neuron-specific isoforms, liprin-α2 and α3, mildly disrupts active zone ultrastructure and vesicle priming in mice ^10^, knocking out all four liprin-α genes in human neurons blocks the recruitment of active zone components and synaptic vesicle ^11^, demonstrating the indispensable role of liprin-α proteins in mammalian presynaptic structure and function. Liprin-α organizes the active zone by interacting with various presynaptic proteins ^6,10,12–16^ (Fig. 1A). Its C-terminal SAM domains associate with presynaptic adhesion molecules, such as LAR-type receptor protein tyrosine phosphatases and neurexins through forming the liprin-α/CASK/neurexin tripartite complex ^11,17–19^. On the other hand, liprin-α employs N-terminal coiled coils to recruit other presynaptic scaffolds, including RIM and ELKS, to synaptic adhesion sites for active zone formation and function ^15,20–23^.

**Figure 1.**
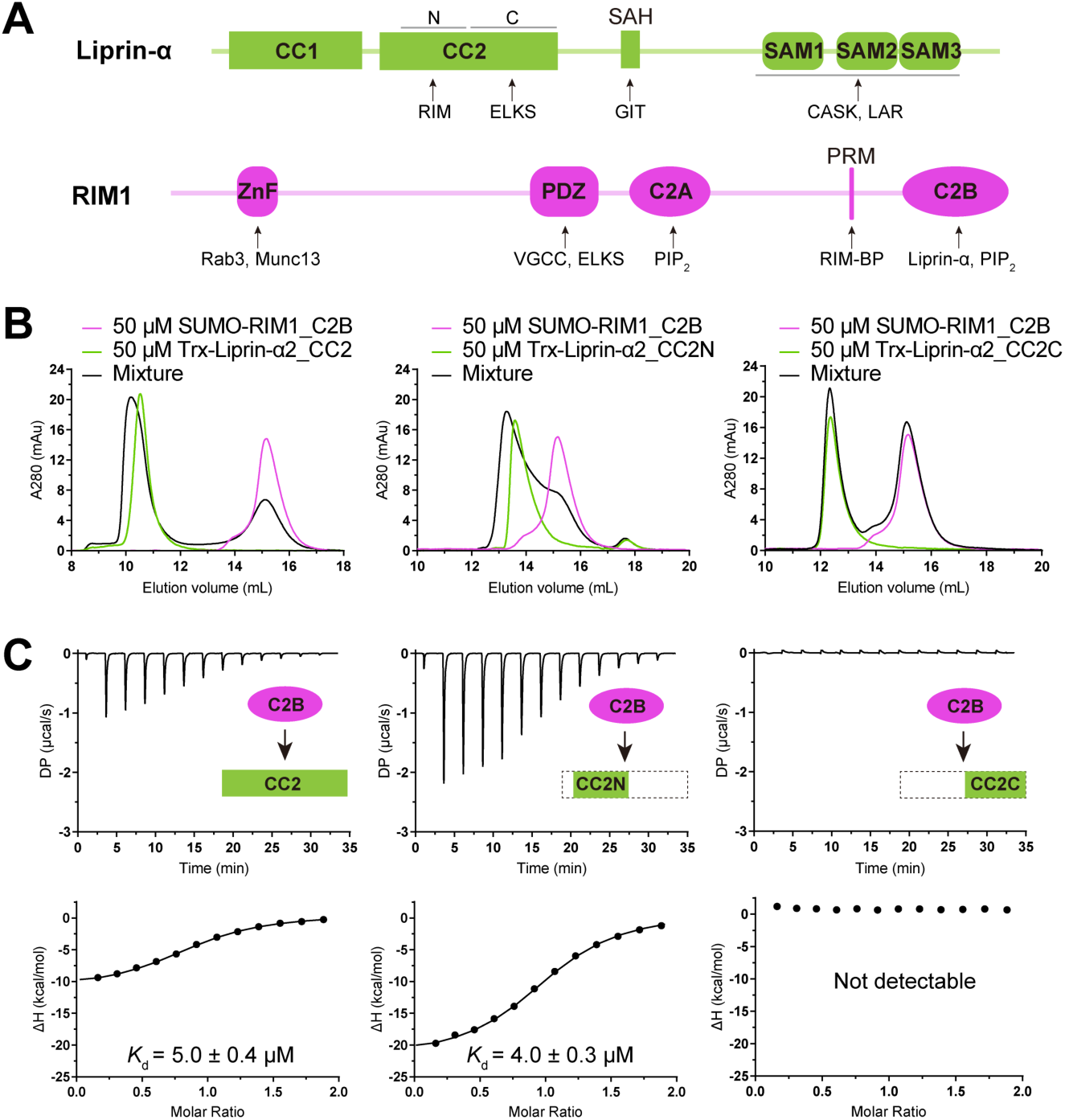
Biochemical characterization of the liprin-α2/RIM1 interaction. **A.** Schematic of liprin-α and RIM1 domain organization, with protein-binding regions indicated. CC, Coiled coil region; SAH, single alpha helix; SAM, sterile alpha motif; ZnF, zinc finger; PRM, proline-rich motif; PDZ, PSD-95/Discs-large/ZO-1 homology; C2, Protein kinase C conserved region 2. **B.** aSEC analysis showing the binding of RIM1_C2B to the N-terminal segment of liprin-α2_CC2. **C.** ITC-based affinity measurement of RIM1_C2B binding to different boundaries of liprin-α2_CC2.

RIM binds to the other four core scaffold proteins and Ca^2+^ channels ^20,24–27^, contributing to active zone assembly. The RIM family in vertebrates has four members ^28^: RIM1 and RIM2 are multidomain-containing proteins with two C-terminal C2 domains (C2A and C2B; Fig. 1A) involved in neurotransmitter release through phospholipid binding ^29^, whereas RIM3 and RIM4 only contain the C2B domain. RIM1 and RIM2 were identified to interact through their C2B domains with the coiled-coil region of liprin-α ^20^. Although the recognized significance of both liprin-α and RIM in active zone formation and function, a molecular understanding of their interaction and its functional consequences remain unknown. Discovery of RIM/RBP condensation indicates the involvement of liquid-liquid phase separation (LLPS) in coupling synaptic vesicles and voltage-gated Ca^2+^ channels (VGCCs) ^30,31^. In agreement with this, liprin-α co-phase separates with ELKS, required for active zone formation ^32^. These findings also raise an intriguing question of how these sophisticated protein assemblies interplay through the liprin-α/RIM interaction.

In this study, we solved the structure of the complex formed by the coiled-coil region of liprin-α2 and the C2B domain of RIM1. The structure uncovers that liprin-α mutations associated with neurodevelopmental diseases block complex formation, supporting the importance of this complex in synapse formation. Further structural analysis reveals a unique binding mode between the coiled-coil dimer and the C2B dimer, which drives a large protein assembly. Through this multivalent binding, liprin-α2 promotes the condensate formation of RIM1, confirming its scaffolding role in early active zone protein recruitment. Using human neurons lacking all liprin-α (liprin-α qKOs), we showed that the liprin-α/RIM interaction is dispensable for synapse formation while required for synaptic transmission and vesicle recruitment. Importantly, our LLPS assays and synaptic analyses indicate that liprin-α2, through its binding to RIM1, not only effectively accumulates RIM1 at the active zone but also promotes VGCC clustering, allowing proximal coupling between VGCC clustering sites and vesicle priming sites for efficient neurotransmitter release. Collectively, our study unveils the presynaptic assembly and regulatory mechanism of active zone machinery via the liprin-α/RIM interaction in an LLPS-dependent manner.

## Results

### Biochemical and crystallographic analyses of the liprin-α2/RIM1 complex

To understand the molecular mechanism governing the liprin-α/RIM interaction, we first identified the minimal segment in liprin-α2 that is sufficient for its binding to the C2B domain of RIM1. As our previous study suggests that liprin-α can form a tripartite complex with RIM and ELKS and a coiled-coil region (CC2; Fig 1A) binds both RIM and ELKS ^23^, we speculated that RIM and ELKS interact with distinct segments of CC2. To validate this, we divided the CC2 region into two parts: the N-terminal half (CC2N) and the C-terminal half (CC2C) (Fig. 1A). The interactions between the two segments and RIM1_C2B were characterized using analytical size-exclusion chromatography (aSEC) and isothermal titration calorimetry (ITC). The results showed that CC2N but not CC2C interacts with RIM1_C2B, with a measured binding affinity of ∼5 μM (Fig. 1B and C). Consistently, the deletion of CC2N from liprin-α2 did not compromise its ability to bind ELKS1, while the deletion of CC2C disrupted the interaction (Fig. S1). These results indicate that liprin-α2 interacts with the C2B domain of RIM1 through its CC2N segment. The differential binding specificity found in the CC2 region provides a mechanism for assembling the liprin-α/RIM/ELKS tripartite complex.

Next, we aimed to solve the structure of the liprin-α2_CC2N/RIM1_C2B complex using X-ray crystallography. Initial attempts to co-crystallize the tag-removed CC2N and C2B fragments resulted in heavy precipitation, preventing crystal formation. To circumvent this issue, we modified our approach by mixing tag-removed liprin-α2_CC2N with SUMO-tagged RIM1_C2B for crystallization. However, this approach also failed to yield any crystals, despite extensive trials, presumably due to the interference of the SUMO tag in protein crystallization. To remove the SUMO tag without inducing severe protein precipitation, we added a trace amount of TEV protease during crystallization. This strategy successfully led to the formation of high-quality crystals, which we used to determine the structure of the liprin-α2_CC2N/RIM1_C2B complex at a resolution of 2.75-Å (Table S1).

### Overall structure of the liprin-α2_CC2N/RIM1_C2B complex

In the complex structure, the CC2N segment forms a dimeric coiled coil that interacts with two RIM1_C2B molecules symmetrically through its N-terminal region, assembling a 2:2 stoichiometric complex (Figs. 2A, S2A, and S2B). Conversely, the C2B domain of RIM1, characterized by a β-sandwich fold, mainly packs with CC2N via a β-sheet composed of strands β-2/3/6/9 (Figs. 2A, 2B, and S2C). The CC2N/C2B interaction is predominantly mediated by polar interactions. Several salt bridges, including R1239^RIM1^-E328/D335^liprin-α2^, R1201^RIM1^-E337^liprin-α2^, and E1198^RIM1^-R346^liprin-α2^, strongly stabilize this interaction (Fig. 2B). Q332^liprin-α2^ and R339^liprin-α2^ also contribute significantly to the binding by forming hydrogen bond networks at the interface (Fig. 2B). In addition to these polar interactions, hydrophobic interactions further strengthen the CC2N/C2B interaction (Fig. 2B). The interface residues in CC2N are highly conserved across liprin-α isoforms in different species (Fig. S2B and D), suggesting that the observed RIM-binding mode is shared by all liprin-α proteins. Conversely, a key interface residue in RIM1, R1201, is not conserved in RIM3 and RIM4 (Fig. 2C). The substitution of R1201 with glutamine in RIM1, to mimic the sequence of RIM4, abolished the CC2N/C2B interaction (Fig. 2D), confirming that the liprin-α/RIM interaction is specific to certain RIM proteins ^29^, including RIM1, RIM2, and their homologs in invertebrates. Likewise, the presence of a glutamine residue at the R1201-corresponding position in the C2A domain of RIM1 explains the selective binding of liprin-α to the C2B domain over C2A (Fig. S2C).

**Figure 2.**
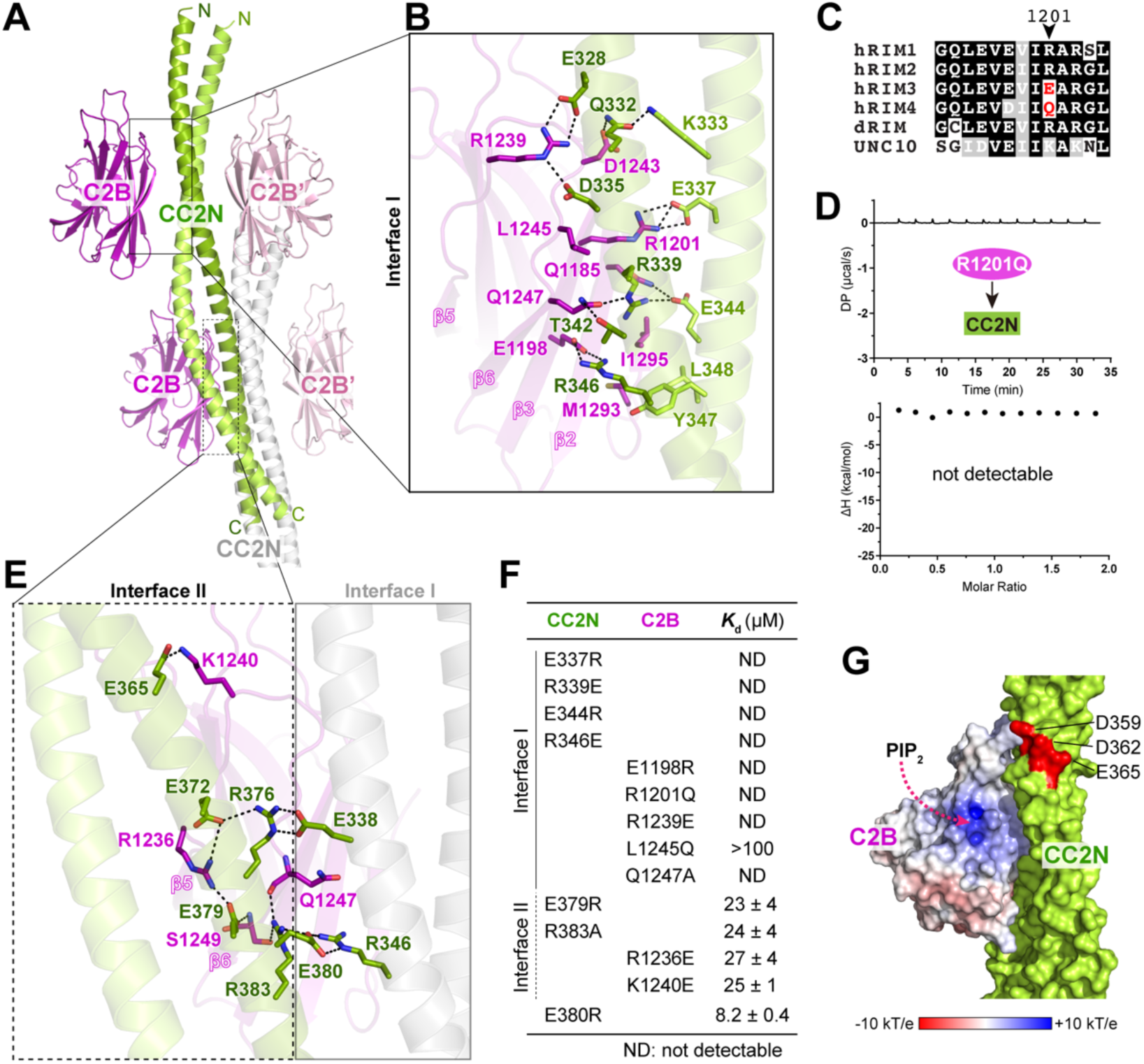
Structural characterization of the liprin-α2_CC2N/RIM1_C2B complex. A. Crystal structure of the liprin-α2_CC2N/RIM1_C2B complex. Two neighboring CC2N coiled coils (colored green and grey, respectively), with four bound C2B molecules (colored magenta), are shown. B. Molecular details of interface I formed between the N-terminal part of CC2N and C2B. Salt bridges and hydrogen bonds are indicated by dashed lines. C. Multisequence alignment of RIM isoforms from various species, showing the sequence variability at the interface residue position corresponding to R1201 in hRIM1. Species abbreviations: ’h’ for human, ’d’ for Drosophila, and UNC10 as the *C.elegans* RIM homolog. D. ITC analysis of the R1201Q RIM1 mutant’s binding to CC2N. E. Molecular details of interface II formed between the C-terminal part of CC2N and C2B. The interconnectivity between interfaces I and II in the complex of one C2B molecule and two CC2N coiled coils is displayed. Salt bridges and hydrogen bonds are indicated by dashed lines. F. Summary of binding affinities between various CC2N and C2B variants, measured by ITC. G. Surface representation of C2B and CC2N, showing the spatial relationship between the PIP_2_-binding site and bound CC2N. Key negatively charged residues on the CC2N structure are highlighted in red.

In addition to the primary interface (interface I) in the formation of the tetrameric complex, the tetramers in the crystal are assembled through a secondary CC2N/C2B interface (interface II) (Fig. 2A). At interface II, the C-terminal part of CC2N interacts with a side face of the β-sandwich fold in RIM1_C2B, mainly through charge-charge interactions and hydrogen bonding (Figs. 2E, S2B, and S3A). ITC-based analysis showed that, while disruptive mutations at interface I abolish the CC2N/C2B interaction, interface II mutations have a milder impact, reducing the binding affinity by ∼5-fold (Figs. 2F, S3B, and S3C). Furthermore, neighboring CC2N coiled coils interact through salt bridges that stabilize both CC2N/C2B interfaces (Figs. 2E and S3A). Specifically, E380 forms salt bridges with R346 in a neighboring CC2N coiled coil, stabilizing the orientation of R346 for its binding to E1198^RIM1^ at interface I (Fig. S3D). The charge-reversed mutation E380R led to a 2-fold decrease in the binding affinity between CC2N and C2B (Figs. 2F and S3E). Together, the crystal structure of the liprin-α2_CC2N/RIM1_C2B complex reveals how the CC2N coiled coil specifically recognizes the C2B domain of RIM1 via two interconnected binding interfaces.

As the C2B domain of RIM1 also binds to phosphatidylinositol 4,5-bisphosphate (PIP_2_) ^29^, we analyzed the potential impact of PIP_2_ on liprin-α’s ability to bind C2B. As shown in Fig. 2G, the putative PIP_2_-binding site on C2B remains fully accessible with bound CC2N. However, when C2B is associated with the PIP_2_-containing membrane, formation of the liprin-α/RIM complex positions a negatively charged patch on the CC2N surface facing the membrane (Fig. 2G). Considering the negatively charged nature of the inner leaflet of the plasma membrane, this spatial arrangement has the potential to generate charge repulsion, thus inhibiting the CC2N/C2B interaction. It suggests that the membrane association of RIM1 in the PIP_2_-enriched compartment may tune its binding to liprin-α.

### Two disease-associated mutations at the CC2N region disrupt the liprin-α/RIM interaction

Many genetic mutations in human liprin-α genes have been linked to neurodevelopmental disorders such as autism, intellectual disability, and epilepsy ^33–36^. Interestingly, several reported missense mutations are located in the CC2N region (Fig. 3A) ^33,35–37^. These mutated sites, including E328^liprin-α2^, A315^liprin-α3^ (corresponding to A350 in liprin-α2), and L330^liprin-α1^ (L348 in liprin-α2), are strictly conserved in the liprin-α family (Fig. 3B). In addition to E328’s critical role in forming salt bridges with RIM1_C2B (Fig. 2B), L348 is directly involved in hydrophobic interactions with M1293 and I1295 in RIM1_C2B (Fig. 2B). A350, although not directly involved in C2B binding, contributes to the coiled-coil formation of CC2 (Figs. 3C and S2B). The charge-reverse mutation E328K disrupts the charge-charge interaction between liprin-α2 and RIM1. To determine the potential consequence of the other two mutations, we performed *in silico* substitutions of the corresponding residues in the complex structure and analyzed the mutated model. As shown in Fig. 3D, the L348F mutation, while retaining hydrophobicity, imposes steric hindrance by introducing a bulkier sidechain, thereby impeding the close contact between CC2N and C2B. However, the A350S mutation, having little impact on the coiled-coil structure of CC2N (Fig. 3E), is unlikely to interfere with the CC2N/C2B interaction.

**Figure 3.**
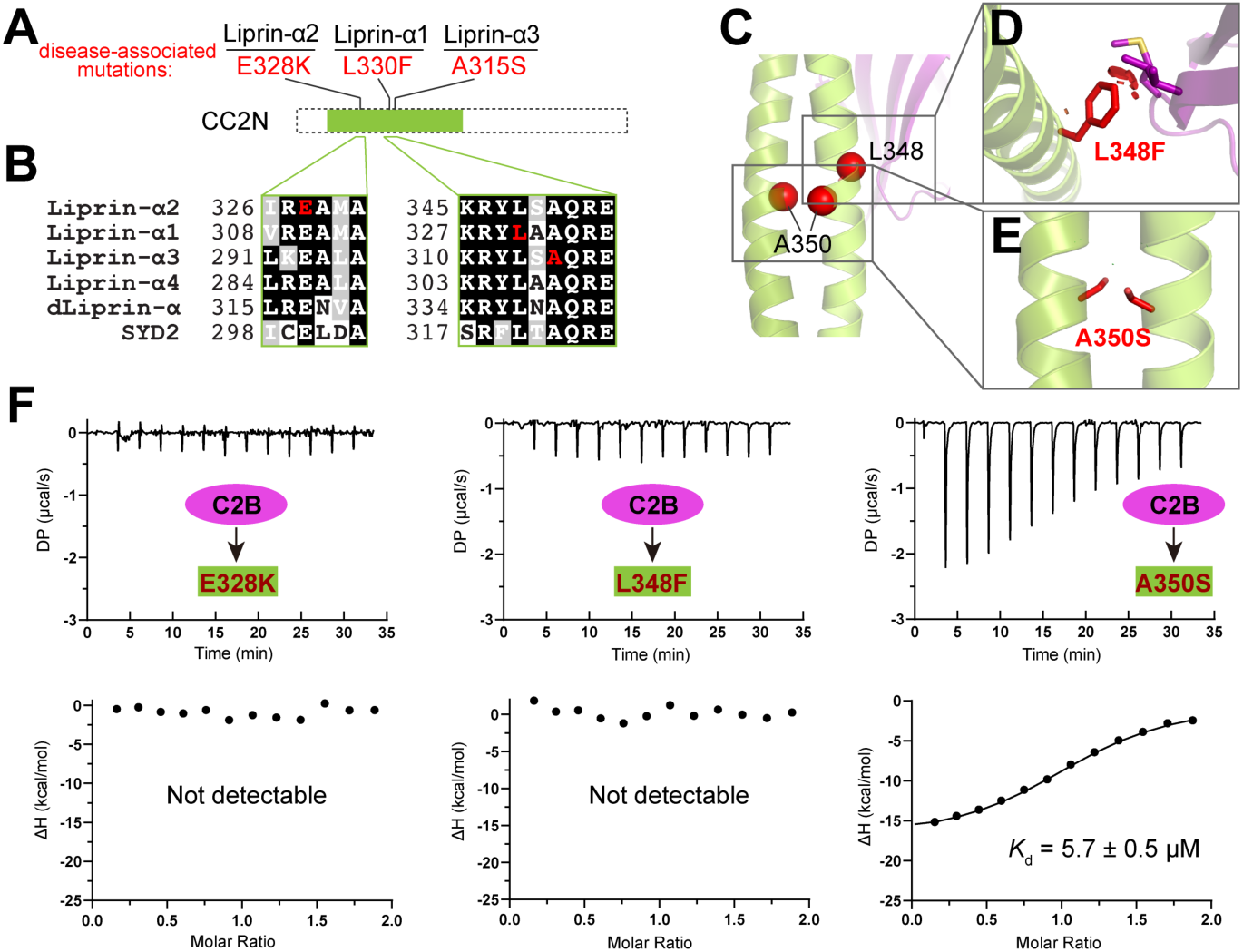
Structural and biochemical analyses of disease-associated mutations on the liprin-α/RIM interaction. **A.** Disease-associated mutations and their positions in the CC2N segment of liprin-α1 (L330, corresponding to L348 in liprin-α2), liprin-α2 (E328), and liprin-α3 (A315, corresponding to A350 in liprin-α2). **B.** Multisequence alignment of liprin-α family members. Residues affected by disease-associated missense variants are marked in red. **C.** Cartoon representation of the liprin-α2_CC2N/RIM1_C2B complex with residues affected by disease-associated missense variants indicated. E328K and L348 are located at interface I, while A350 contributes to the coiled-coil formation of CC2N. **D.** Structural analysis of the L348F mutation showing steric hindrance caused by the mutated sidechain upon the complex formation. Atomic clashes are indicated by red cylinders. **E.** Structural analysis of the A350S mutation in the context of the coiled-coil structure, revealing no disruption. **F.** ITC-based analyses of the interactions between C2B and the E328K, L348F, and A350S mutants of CC2N.

To further evaluate the mutational effects on the liprin-α/RIM interaction, we introduced E328K, L348F, and A350S mutations to the CC2N construct and measured the binding affinities of the CC2N mutants to RIM1_C2B. Consistent with our structural analysis, the E328K and L348F mutations eliminate the CC2N/C2B interaction, while the A350S mutation had minimal impact on binding affinity (Fig. 3F). Considering that the L330F mutation in liprin-α1 was identified in patients with autism ^35^, our results suggest that this mutation may impair synapse development by interfering with the binding of liprin-α1 to RIM proteins. Nevertheless, given the poorly defined function of liprin-α1 at the presynapse, whether the L330F mutation contributes to neurodevelopmental defects through the disruption of liprin-α1/RIM interaction requires further investigation.

### Liprin-α2 and RIM1 form a large complex through multivalent binding

Although RIM1_C2B was purified as a monomer, it forms a homodimer in our crystal structure, a feature also observed in the apo C2B structure of RIM1 ^38^ (Figs. 4A and S4A). This dimerization tendency was confirmed in solution, as increasing concentrations of RIM1_C2B led to a corresponding increase in dimer formation (Figs. 4B and S4B). Considering the C2B dimerization along with the CC2N/C2B interactions, we propose that liprin-α2 and RIM1 may assemble into a large complex through a network of intermolecular interactions revealed in our crystal structure (Figs. 4C and S4A). This hypothesis is supported by aSEC analysis of a 500 μM liprin-α2_CC2/RIM1_C2B mixture, which showed the formation of protein assemblies with molecular weights even larger than 500 kDa (Fig. 4D). In contrast, either CC2 or C2B alone maintains their dimeric state even at a concentration of 500 μM (Fig. S4B and C). These results suggest the involvement of multivalent interactions in assembling the liprin-α2/RIM1 complex. However, given the high protein concentrations (sub-mM) used in our crystallographic and aSEC analyses, it prompts an important question of whether the observed multivalent interactions could occur in the context of presynaptic assembly.

**Figure 4.**
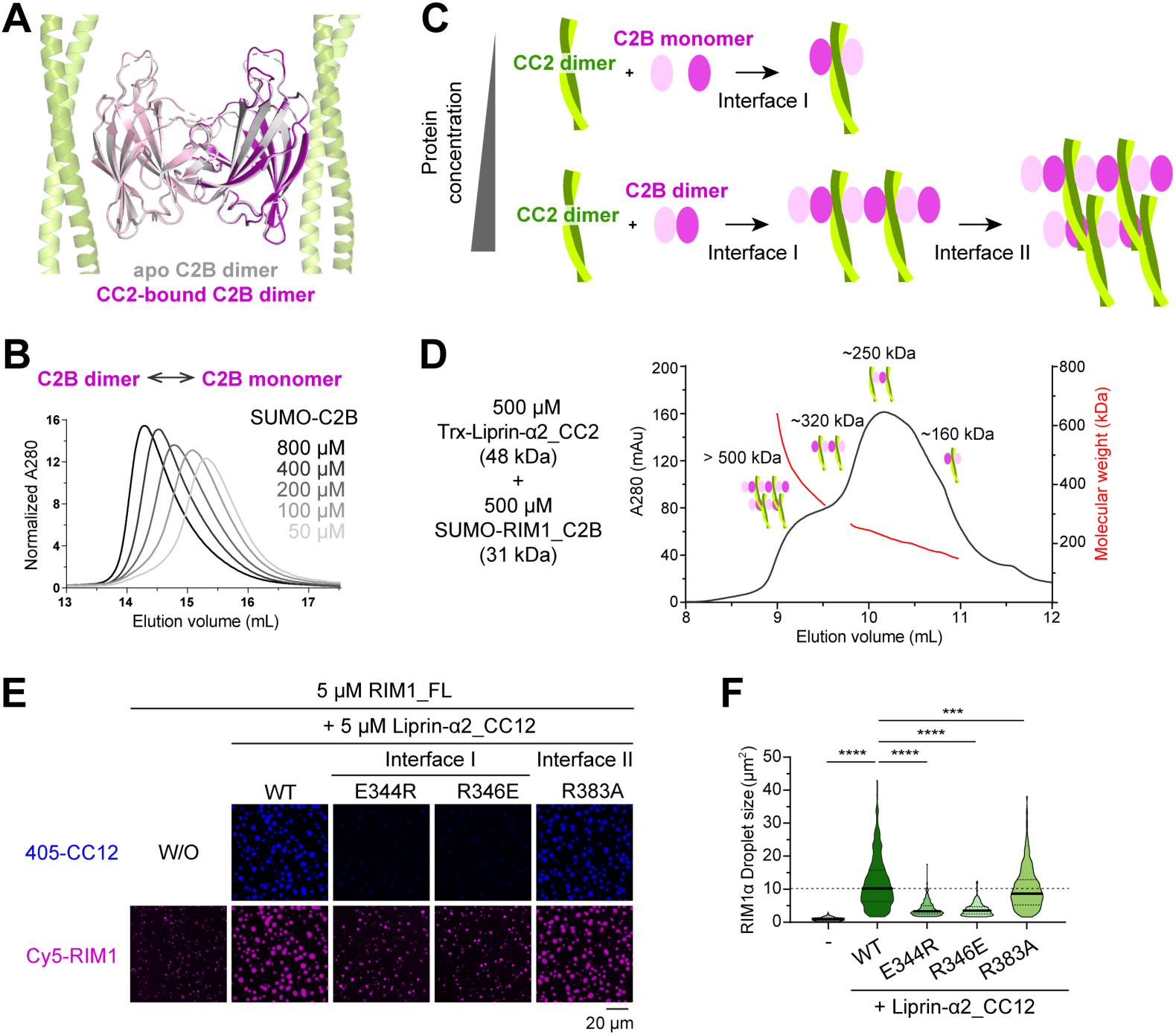
Unique assembly mode of the liprin-α2/RIM1 complex. **A.** Structural superimposition of the RIM1_C2B dimer from two crystal structures. The overall RMSD of these two dimeric C2B structures is 0.4 Å. **B.** Concentration-dependent dimerization of RIM1_C2B in solution. **C.** Schematic representation of the liprin-α2_CC2N/RIM1_C2B complex assembly modes. Under low-concentration conditions, a 2:2 heterodimer is formed, whereas high-concentration conditions lead to the assembly of a large complex through multiple intermolecular interactions. **D.** aSEC analysis coupled with multi-angle static light scattering (MALS), showing the formation of large CC2N and C2B assemblies. **E.** *In vitro* LLPS assays showing the CC2N/C2B assembly in promoting RIM1 LLPS. **F.** Quantification analysis of RIM1 droplet sizes presented in panel **E**. Droplets from 8 different views are quantified, and all data are represented as means ± SD. The unpaired Student’s *t* test analysis was used to define a statistically significant difference (****, p < 0.0001; ***, p < 0.001).

Recent studies on active zone proteins, including liprin-α, RIM, and ELKS, reveal their pronounced propensity for LLPS ^23,30,32,39^. Through LLPS, these proteins can concentrate within condensates ranging from sub-mM to mM levels ^23,30^. Given the well-established significance of multivalent binding in mediating LLPS ^40,41^, it is plausible that liprin-α2 and RIM1 may coalesce into a co-condensate, presumably facilitated by the multivalent interactions identified in our crystal structure. To explore the potential role of the CC2N/C2B interaction in the co-condensation of liprin-α2 and RIM1, we performed *in vitro* LLPS assays with a purified N-terminal segment of liprin-α2 (liprin-α2_CC12) and full-length RIM1. Notably, liprin-α2_CC12, containing both the CC1 and CC2 regions, has been shown to promote the LLPS of ELKS ^23^. Indeed, compared to the condensate formed by RIM1 alone, the addition of liprin-α2_CC12 robustly enlarged the RIM1 droplet size (Fig. 4E and F). In contrast, the E334R and R346E mutations at interface I of CC12 diminished the promotive effect, confirming the importance of the CC2N/C2B interaction in promoting RIM1 LLPS. Additionally, the R383A mutation at interface II of CC12 modestly attenuated the enhancing effect on droplet size (Fig. 4E and F), consistent with the milder impact of interface II on the disruption of the CC2N/C2B interaction (Fig. 2F). Similarly, our *in vitro* sedimentation-based assay confirmed the CC12-mediated promotion effect of RIM1 LLPS (Fig. S4D and E). Altogether, our structural and biochemical analyses highlight the role of the CC2N/C2B interaction in assembling liprin-α2 and RIM1 into condensates, offering insights into the molecular mechanism underlying the dynamic assembly of the presynaptic active zone.

### The CC2N/C2B interaction is dispensable for synaptic formation

To study the functional role of the liprin-α2/RIM1 complex in active zone assembly and function, we employed pluripotent stem-cell-derived human neurons lacking all liprin-α proteins (liprin-α qKO neurons) ^11^, in which the assembly of active zones and the recruitment of synaptic vesicles to nascent terminals is completely blocked. By re-expressing wild-type (WT) liprin-α2 or its mutants, including the CC2N deletion (ΔCC2N) and point mutations E344R and R346E, we can specifically analyze the CC2N/C2B interaction’s effects on synaptic defects in liprin-α qKO neurons, as these mutants disrupt RIM1 binding while retaining ELKS1 interaction (Fig. S5A and B). To ensure comparable expression levels of liprin-α2 variants, an approach of lentivirus transduction was used (Fig. S5C).

Quantitative analysis of synapses using synapsin and MAP staining for pre-and post-synaptic compartments, respectively, showed that the deletion of all liprin-α isoforms almost eliminated synaptic puncta as we reported previously ^11^, but re-expression of either liprin-α2 WT or RIM-binding-deficient mutants comparably restored synapsin puncta signals in liprin-α qKO neurons (Fig. S6A and B). Next, we compared synaptic levels of active zone proteins in liprin-α qKO neurons rescued with either liprin-α2 or its mutants using confocal microscopy. No significant differences were found in the overall synaptic levels of these active zone proteins between the rescues with the WT and mutant constructs (Fig. S6C and D). These findings suggest that the disruption of the liprin-α2/RIM1 complex has no major effects on active zone assembly, and the CC2N/C2B interaction plays a dispensable role in synaptic formation.

### The CC2N/C2B interaction is critical for synaptic transmission

To further assess the significance of the liprin-α/RIM complex on synaptic function, we performed direct measurements of miniature excitatory postsynaptic currents (mEPSCs) in liprin-α qKO neurons. The absence of all four liprin-α isoforms nearly abolished spontaneous synaptic transmission (Fig. 5A-C), consistent with our previous findings ^11^. We then rescued the synaptic deficit in liprin-α qKO neurons by re-expressing liprin-α2 or its mutants. As shown in Fig. 5A-C, WT liprin-α2 re-expression substantially restored mEPSCs, whereas the mutants were significantly less effective. Considering the specific disruption of the liprin-α/RIM interaction by the mutations, these results strongly support the role of liprin-α in conjugation with RIM proteins via the CC2N/C2B interaction to regulate synaptic vesicle release and neurotransmission.

**Figure 5.**
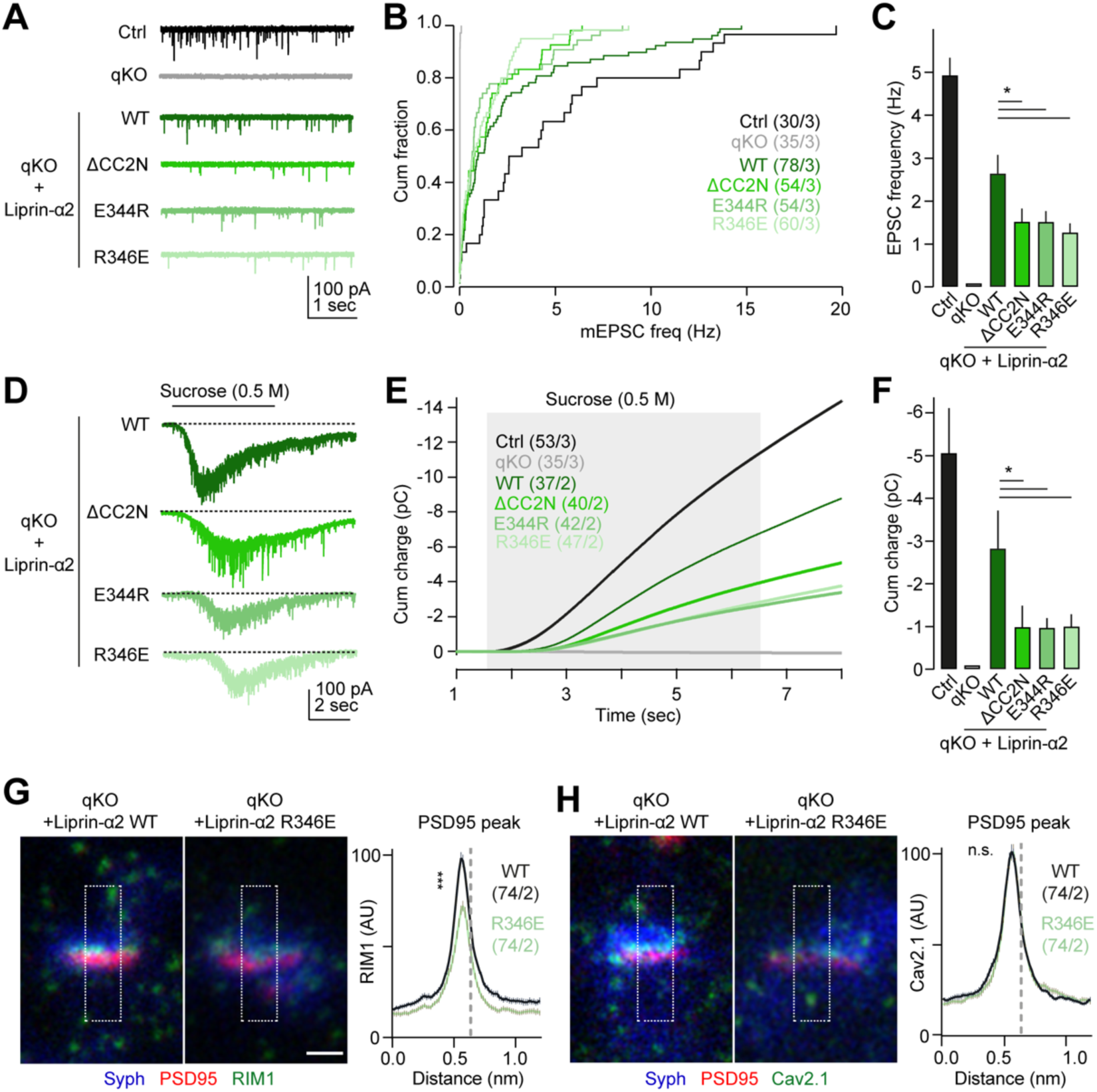
Liprin-α2/RIM1 complex controls synaptic function. **A.** Representative traces of spontaneous glutamatergic transmission, measured as miniature excitatory postsynaptic currents (mEPSCs) in Control (Ctrl) neurons, liprin-α quadruple knockout (qKO), and qKO neurons rescued with liprin-α2 WT or RIM-binding-deficient mutants (ΔCC2N, E344R, and R346E). Recordings were performed at a holding potential of - 70 mV and in the presence of 0.5 μM tetrodotoxin (TTX). **B.** Cumulative distributions of EPSC frequency under the conditions indicated in panel **A**. **C.** Statistical analyses of **B** showing the impact of liprin-α2 mutations on synaptic transmission. The number of cells/batches analyzed for each condition was indicated in panel **B**. Data represented as means ± SEM; * p < 0.05. **D.** Representative traces showing the response to hyperosmotic sucrose in indicated conditions. Neurons were challenged with 0.5 M sucrose for 5 seconds (shaded area in panel **E**) using a flow pipe placed in close proximity (near 100 μm) to the recorded cells. **E.** Integrated responses (EPSC charge) to hypertonic sucrose application in indicated conditions. **F.** Statistical analyses of **E** showing the impact of liprin-α2 mutations on the size of the RRP. The number of cells/batches analyzed for each condition was indicated in panel **E**. Data represented as means ± SEM; * p < 0.05. **G, H.** Subsynaptic imaging and summary plots of RIM1 (**G**) and Cav2.1 (**H**) intensity profiles in liprin-α qKO neurons rescued with liprin-α2 WT (black) or R346E mutant (light green). The relative peak of the PSD95 signal is indicated by the vertical dotted line. Scale bar: 250 nm. Number of profiles/batches analyzed: liprin-α2 WT:74/2; liprin-α2 R346E:74/2. Data represented as means ± SEM; n.s. non-significant, *** p < 0.001.

Next, we explored whether the liprin-α/RIM complex influences synaptic transmission by regulating primed synaptic vesicles by analyzing the size of the readily releasable pool (RRP) of synaptic vesicles, using hyperosmotic sucrose treatment as described earlier ^42^. In liprin-α qKO neurons, sucrose responses were eliminated but were readily rescued by liprin-α2 WT re-expression (Fig. 5D-F). In contrast, RIM-binding-deficient mutants only partially rescued sucrose responses, which were significantly smaller than those triggered by the WT rescue (Fig. 5D-F). It suggests that the liprin-α/RIM interaction is important in maintaining the RRP size. Together, our structural and functional results demonstrate that the liprin-α/RIM complex, assembled by the CC2N/C2B interaction, controls the synaptic transmission, at least in part, by regulating the number of primed synaptic vesicles in nerve terminals.

### Liprin-α2 facilitates the presynaptic accumulation of RIM1 through the CC2N/C2B interaction

Given the essential role of RIM1 in vesicle priming at the active zone ^43,44^, the liprin-α/RIM1 complex may regulate RIM1 levels at the active zone, which in turn could influence synaptic vesicle priming and release. To test this possibility, we analyzed the presynaptic level of RIM1 using STED super-resolution microscopy (Fig. S6E). We found a significant reduction of RIM1 signals at the presynaptic termini under the R346E rescue condition, compared to the WT liprin-α2 (Fig. 5G). This finding, coupled with the role of liprin-α2 in promoting RIM1 condensate formation through the CC2N/C2B interaction (Fig. 4E and F), suggests that liprin-α may effectively accumulate RIM1 at the active zone by forming the liprin-α/RIM1 complex. As RIM1 may also be involved in recruiting VGCCs to the active zone ^26,45^, we compared presynaptic P/Q-type VGCC α1 subunit (CaV2.1) levels between the WT and R346E rescue conditions. However, no significant alternation in CaV2.1 levels was detected (Fig. 5H), suggesting that while the liprin-α/RIM interaction helps recruiting RIM1 to nerve terminals, it does not control the overall density of VGCCs at the active zone. Nevertheless, potential nanoscale changes in Ca^2+^channel positioning upon liprin-α/RIM interaction disruption cannot be readily ruled out. Indeed, previous studies at the calyx of Held synapses have clearly shown changes in presynaptic VGCC nano-organization without altering the overall density of these channels in nerve terminals ^46^.

### The liprin-α/RIM complex controls the clustering of Ca^2+^ channels through mesoscale interactions among ELKS1 and RIM1 condensates

Liprin-α, through its CC2 region, assembles ELKS and RIM proteins (Fig. 6A), which contribute to the nano-scale organization of presynaptic VGCCs ^26,47^. Therefore, we hypothesize that the liprin-α/RIM complex may cooperate with ELKS to regulate the local distribution of VGCCs, without affecting the overall level of Ca^2+^channels at the active zone. Intriguingly, both ELKS1 and RIM1 can form condensates via LLPS, regulated by liprin-α and RBP, respectively ^23,30^. To explore the relationship between the ELKS1 and RIM1 condensates, we prepared these condensates using purified full-length proteins. To enhance RIM1 LLPS, the RBP2_(SH3)_3_ fragment was added (Fig. 6A), as reported previously ^30^. Without liprin-α2_CC12, the ELKS1 and RIM1 condensates merge to form co-phase droplets (Fig. 6B). However, the presence of liprin-α2_CC12 prevents co-condensation, with the droplets of ELKS1 and RIM1 become largely immiscible (Fig. 6C). Consistent with our previous findings using ELKS2 fragments ^23^, ELKS1/liprin-α2_CC12 was found to enrich at the periphery of RIM1 droplets (Fig. 6C, box ‘a’). Conversely, no RIM1 accumulation was observed in ELKS1 droplets (Fig. 6C, box ‘b’), although both liprin-α2_CC12 and ELKS1 can interact with RIM1 (Fig. 6A).

**Figure 6.**
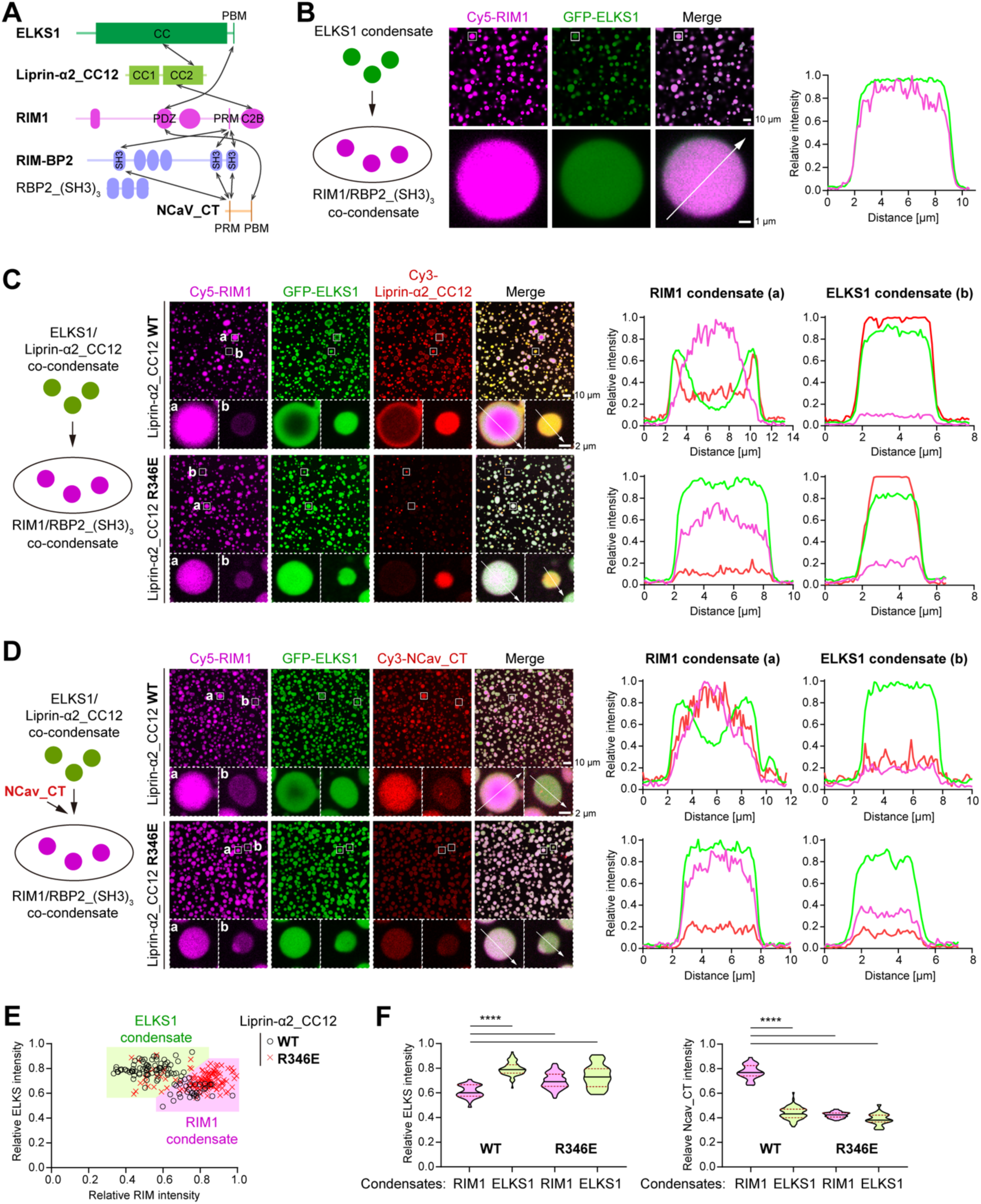
The liprin-α/RIM interaction modulates the distribution of ELKS1 and NCav_CT in RIM1 condensates. **A.** Schematic diagram of the interaction network among active zone proteins. Interactions are indicated by the double-headed arrows. **B.** Confocal imaging analysis of the LLPS mixture containing the ELKS1 condensate and the RIM1/RBP2_(SH3)_3_ condensate. A magnified view of a representative droplet was displayed below, with a line analysis of fluorescence signal intensities along the indicated dashed line. The concentration of each protein was 5 μM. **C.** Confocal imaging analysis of the LLPS mixture containing the ELKS1/liprin-α2_CC12 (WT or R346E) condensate and the RIM1/RBP2_(SH3)_3_ condensate. Magnified views of representative RIM1 (a) and ELKS1 (b) droplets were displayed below, with line analyses of fluorescence signal intensities along the indicated dashed lines. The concentration of each protein was 5 μM. **D.** Confocal imaging analysis of the LLPS mixture containing the ELKS1/liprin-α2_CC12 (WT or R346E) condensate, the RIM1/RBP2_(SH3)_3_ condensate, and NCav_CT. Magnified views of representative RIM1 (a) and ELKS1 (b) droplets were displayed below, with line analyses of fluorescence signal intensities along the indicated dashed lines. The concentration of each protein was 5 μM. **E.** Plot analyses of the intensity relationship between ELKS1 and RIM1 fluorescence signals in the condensates shown in panel **D**. The intensity of ∼100 droplets in the view was quantified and normalized. The relative intensity ratio of RIM1/ELKS1 >1 was defined as RIM1 condensate, while the ratio <1 was defined as ELKS1 condensate. **F.** Quantitative analyses of RIM1, ELKS1, and NCav_CT fluorescence intensities in the ELKS1 and RIM1 condensates. Data represented as means ± SD. The unpaired Student’s *t* test analysis was used to define a statistically significant difference (****, p < 0.0001).

The disruption of the liprin-α/RIM interaction by introducing R346E into liprin-α2_CC12 eliminated the surface coating of CC12 on RIM1 droplets (Fig. 6C), indicating that the liprin-α/RIM interaction is essential for recruiting CC12 onto RIM1 condensates. However, the absence of liprin-α2_CC12 from RIM droplets led to an increased accumulation of ELKS1 within RIM1 droplets, aligning with the formation of ELKS1/RIM1 co-phase droplets in the absence of CC12 (Fig. 6B). These observations suggest that the liprin-α/RIM interaction limits the accumulation of ELKS1 within the RIM1 condensate. Additionally, without the addition of RBP2_(SH3)_3_, RIM1, ELKS1, and liprin-α2_CC12 form co-phase droplets (Fig. S7A), indicating that RBP2 also contributes to the immiscibility between the RIM1 and ELKS1 condensates when liprin-α2_CC12 is present. Consistently, liprin-α2_CC12 is weakly enriched on the periphery of RIM1/RBP2_(SH3)_3_ co-condensates (Fig. S7B), compared to its co-condensation formation with RIM1 alone (Fig. 4E). Thus, the peripheral enrichment of liprin-α2_CC12 changes the surface property of the RIM1 condensates, which potentially hinders the diffusion of ELKS1 molecules into condensates. This hypothesis is supported by fluorescence recovery after photobleaching (FRAP) analyses, which show a decrease in the dynamic properties of RIM1 condensates in the presence of liprin-α2_CC12 compared to RBP2_(SH3)_3_ (Fig. S7C).

The RIM1 condensate is known to enrich the cytoplasmic tail of presynaptic Ca^2+^ channels ^23,30^, which interacts with the PDZ domain of RIM (Fig. 6A) ^26^. As ELKS interacts with the PDZ domain of RIM with a much higher binding affinity ^24,25,30,48^, the accumulation of ELKS1 in RIM1 condensates may prevent the enrichment of VGCCs through binding competition. Indeed, the addition of the cytoplasmic tail of the N-type VGCC α1 subunit (NCav_CT) to the mixture of ELKS1 and RIM1 condensates resulted in its accumulation in RIM1 condensates (Fig. 6D). However, disrupting the liprin-α/RIM interaction decreased the overall level of NCav_CT in RIM1 condensates, presumably due to the accumulation of ELKS1 in RIM1 condensates (Fig. 6D). By classifying droplets based on their fluorescence intensity relationship (Fig. 6E), we quantitatively compared the ELKS1 and NCav_CT intensities in the two classes of condensates under different conditions (Figs. 6F and S7D). The results confirmed that disrupting the liprin-α/RIM interaction significantly increases the distribution of ELKS1 in RIM1 condensates (Fig. 6F, left panel), which in turn dramatically reduces the distribution of NCav_CT in RIM1 condensates to a level comparable to that in ELKS1 condensates (Fig. 6F, right panel).

Taken together, our LLPS analyses demonstrate that the liprin-α/RIM complex plays a regulatory role in the distribution of both ELKS1 and VGCCs in the RIM1 condensate. Considering the essential role of VGCC nano-scale clustering in efficient vesicle release, liprin-α may control the interplay between RIM, ELKS, and VGCCs in synaptic transmission by regulating mesoscale protein-protein interactions in the condensed phase.

### The liprin-α/RIM complex promotes efficient coupling between presynaptic Ca^2+^ channels and primed synaptic vesicles

We then directly assessed if liprin-α/RIM complex, by regulating the accumulation of ELKS1 in RIM1 condensates, can control the fine-scale localization of VGCCs within the active zone and thereby regulate the nanodomain coupling between VGCCs and primed synaptic vesicles. For this, we used a channel-rhodopsin-assisted approach to assess evoked synaptic transmission in liprin-α qKO neurons rescued with either liprin-α2 WT or R346E (Fig. 7A). We loaded nerve terminals with the ‘slow’ Ca^2+^-chelator EGTA-AM, which selectively chelates diffusing Ca^2+^ ions not involved in nanodomain coupling between Ca^2+^ channels and the release machinery (Fig. 7B). We reasoned that if the liprin-α/RIM complex is critical for this short-distance coupling, evoked release in the R346E rescue would be more sensitive to EGTA-AM than that in the WT rescue. Indeed, we observed that EGTA-AM more effectively blocked evoked release in liprin-α qKO neurons rescued with R346E, compared to those rescued with WT construct (Fig. 7C and D), suggesting an increased distance between Ca^2+^ channels and Ca^2+^ sensors in the release machinery upon disruption of the liprin-α/RIM interaction. Control treatment with vehicle (DMSO) showed no significant differences in evoked release (Fig. 7C and D). Altogether, these results indicate that the liprin-α/RIM complex is crucial for localizing Ca^2+^channels in close proximity to primed synaptic vesicles within the active zone and to ensure tight coupling between presynaptic action potentials and neurotransmitter release. These finding also provides a plausible mechanism for liprin-α/RIM complexes in the regulation of the nanoscale VGCC clustering, while having a negligible impact on its overall presynaptic levels (Fig. 5H) that may involve alternative interactions between VGCC and other active zone components.

**Figure 7.**
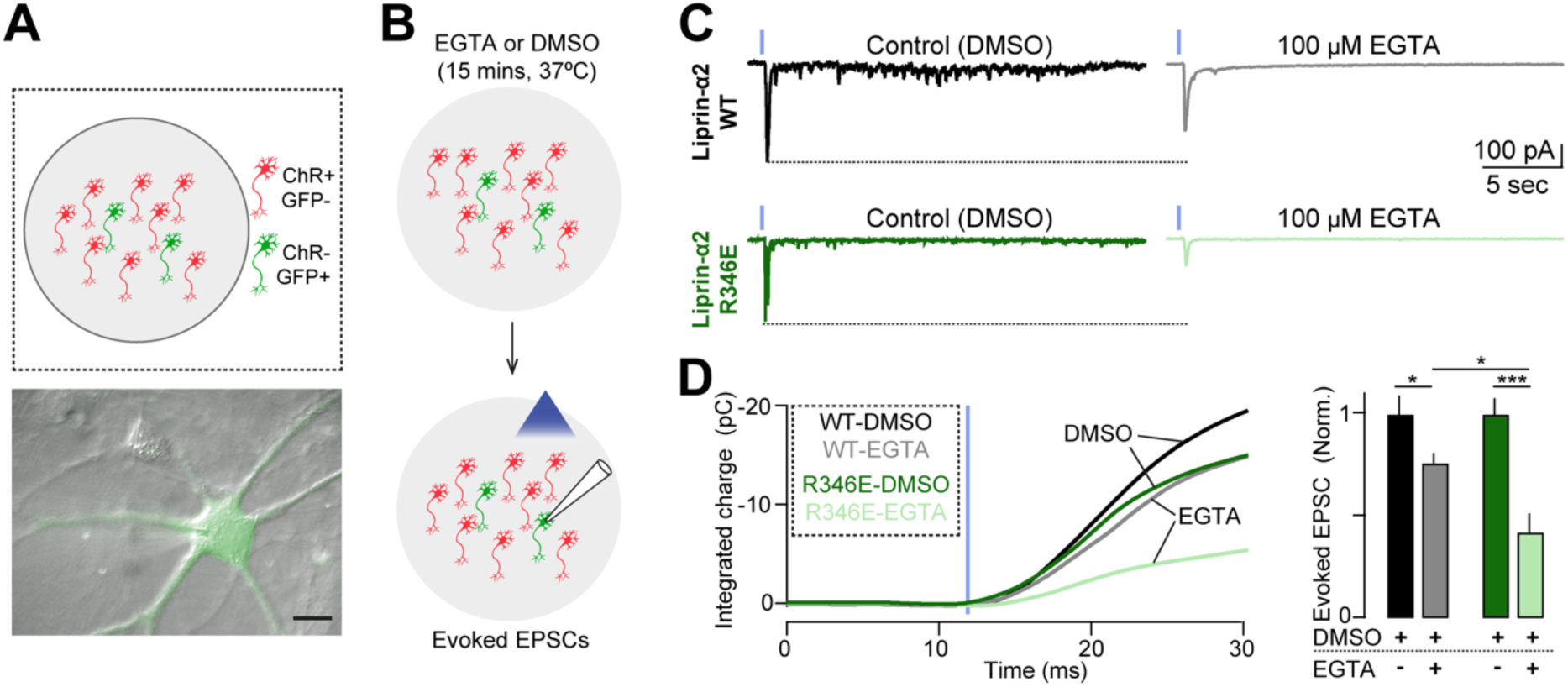
Liprin-α/RIM complexes couple presynaptic Ca^2+^ channels with primed synaptic vesicles. **A.** Top. Schematic representation of the experimental approach for recording channel-rhodopsin-assisted evoked EPSCs in liprin-α qKO neurons rescued with liprin-α2 WT or R346E mutants. Channelrhodospin (ChR^+^/GFP^-^)- and GFP(ChR^-^/GFP^+^)-expressing neurons were seeded at 4/1 ratios, respectively. Bottom: a montage of fluorescent and DIC images showing the patch clamp pipette approaching a ChR^-^/GFP^+^ neuron. Scale bar: 15 μm. **B.** Neurons were treated with 100 μM EGTA-AM (dissolved in DMSO) or DMSO (vehicle) as a control. Evoked excitatory postsynaptic currents were triggered using blue light to activate channel-rhodopsin expressed presynaptically and monitored postsynaptically using whole-cell voltage-clamp recordings. **C.** Representative recordings of evoked EPSCs in liprin-α qKO neurons rescued with liprin-α2 WT (top) or R346E (bottom) constructs. The averaged responses of consecutive 5 trials (at 60 seconds intervals) are shown, in neurons treated with DMSO (left) or EGTA (right). The timing of light activation was indicated by vertical blue ticks. **D.** Summary graphs of evoked glutamatergic transmission in liprin-α qKO neurons rescued with liprin-α2 WT (black) or R346E (green) constructs, under control (DMSO, dark) conditions or under EGTA (light) treatment. Left, averaged integrated responses. Right, responses normalized by the peak of evoked currents. Number of cells/batches analyzed: L2-WT DMSO: 53/2; L2-WT EGTA: 55/2; L2-R346E DMSO: 64/2; L2-R346E EGTA: 49/2. Data represented as means ± SEM; * p < 0.05; *** p < 0.001.

## Discussion

Our study unveils the unique assembly mechanism of the liprin-α/RIM complex, a critical yet less characterized interaction in the presynaptic active zone. Through a combination of structural biology and biochemical approaches, we have elucidated the sophisticated molecular interactions that govern the formation of this complex. Remarkably, our discovery of the liprin-α/RIM interaction’s role in orchestrating the interplay of the RIM1 and ELKS1 condensates provides fresh insights into the nano-scale organization of the active zone. Supported by synaptic analyses in liprin-α qKO neurons with rescue assays, the functional significance of this interaction extends to the modulation of synaptic vesicle release, specifically through its impact on RIM distribution and VGCC clustering. Thus, our work deepens the understanding of the active zone’s assembly mediated by liprin-α and RIM, as well as the dynamic coupling between protein machinery for vesicle priming and release in synaptic transmission.

Our results indicate that liprin-αs, via direct interactions with RIM, can dynamically control two essential functions of the active zone, namely synaptic vesicle priming and subsynaptic distribution of VGCCs, although via different mechanisms. The defects in vesicle priming observed upon disruption of liprin-α/RIM complexes (Fig. 5) can be readily explained by a concomitant reduction in the levels of RIM1 at the active zone, which in turn prime synaptic vesicles via direct interactions with Munc13. In contrast, the defects in VGCC nanoscale localization (Figs. 6 and 7) can be explained in part by the reduction of active zone RIM but perhaps more importantly by a redistribution of ELKS1 in RIM1 condensates, which can outcompete and displace VGCC bound to RIM PDZ domains, resulting in a dramatic reduction of VGCC levels in RIM1 condensates. Thus, our results align with the idea that liprin-αs act as master organizers of presynaptic assembly, regulating the nano-organization of the active zone functions via a multitiered interaction network. At the core, liprin-αs can dynamically and directly interact with RIM and ELKS in a single protein complex, which further coordinates other presynaptic components such as Munc13, RBP, and calcium channels.

Our findings support the involvement of LLPS in the dynamic organization of presynaptic active zones. Emerging evidence has highlighted the nanoscale clustering of RIM ^49^, neurexin ^50^, Munc13 ^51^, and Ca^2+^ channels ^52,53^ at subregions of the active zones. Considering the active zone’s capacity to adapt to various stimuli, LLPS likely facilitates the rapid reorganization of these presynaptic nanoclusters, enabling swift adjustments to synaptic activity. Mesoscale interactions between membrane-associated condensates (e.g., ELKS1 and RIM1 condensates) may contribute to such a dynamic organization of these presynaptic nanoclusters, providing mechanistic insights into the adaptable assembly of the active zone. In this framework, liprin-α emerges as a central regulatory hub, controlling the interplay between active zone proteins in the context of LLPS. Specifically, the liprin-α/RIM interaction not only promotes the recruitment of RIM1 to the active zone (Fig. 5G) but also restricts the enrichment of ELKS1 in RIM1 condensates, maintaining the required VGCC clustering (Fig. 6C-F) for synaptic transmission. Thus, the observed immiscibility between the RIM1 and ELKS1 condensates may be necessary for the nano-scale organization at the active zone. This may ensure that highly condensed active zone proteins maintain their different localization and clustering, allowing them to form distinct functional assemblies at different sites during neurotransmitter release.

The intricate interaction network among active zone proteins is likely to provide a critical layer of functional redundancy that ensures the robustness and adaptability of synaptic connections ^1,3^. For instance, the disruption of the liprin-α/RIM interaction does not completely block the recruitment of RIM1 to the active zone (Fig. 5A and B), presumably due to compensation by the binding of RIM to RBP and ELKS (Fig. 6A). This compensatory effect explains the significant yet not profound impairment in synaptic function (Fig. 5). Nevertheless, given the role of liprin-α in early synaptic development ^11,14,22,54^, the liprin-α/RIM interaction could be critical for the proper assembly of synapses. The compromised RIM protein recruitment by disruptive mutations in liprin-α may still result in a deficit in synaptic homeostasis, contributing to pathological conditions like neurodevelopmental diseases associated with liprin-α proteins ^33,34^.

Interestingly, Rabphilin-3A and synaptotagmin, known for their roles in the docking and fusion of synaptic vesicles, also utilize their C2B domains to interact with the coiled-coil structure of the SNARE complex through multiple interfaces ^55,56^. This binding mode similarity with the liprin-α/RIM complex suggests a general mechanism by which multi-interface binding can modulate the assembly and function of protein complexes in response to synaptic activity. In addition, the C2B domains of Rabphilin-3A and synaptotagmin are also involved in lipid binding ^57^. The binding of lipids to the C2B domain could serve as a molecular switch that influences the conformation, localization, or interaction partners of these C2B-containing proteins (Fig. 2G), which have been extensively studied in synaptotagmin ^58–60^. Given the similar role of the C2B domain of RIM1 in vesicle release ^29^, it is likely that the coupling mechanism between membrane and liprin-α interactions of the C2B domain in RIMs also contributes to neurotransmitter release, which is compelling to be further investigated.

## Acknowledgments

We thank the assistance of Southern University of Science and Technology (SUSTech) Core Research Facilities and Shanghai Synchrotron Radiation Facility (beamlines 17U1, 18U1, and 19U1). We thank Prof. Pilong Li for the pETL7 vector. This work was supported by Key-Area Research and Development Program of Guangdong Province (Grant No. 2023B0303010001), Shenzhen Science and Technology Program (RCJC20210609104333007 to Z.W.), the National Natural Science Foundation of China (32170697 to C.Y. and 32371009 to X.X.), Shenzhen-Hong Kong Institute of Brain Science, Shenzhen Fundamental Research Institutions (2023SHIBS0002), Shenzhen Key Laboratory of Biomolecular Assembling and Regulation (ZDSYS20220402111000001), the Chica and Heinz Schaller Foundation (C.A.), Brain and Behavior Research Foundation (C.A.), the Deutsche Forschungsgemeinschaft (DFG SFB1158-SO2, C.A.), the Fritz Thyssen Foundation (grant 10.21.0.019MN, C.A.), the Swedish Research Council (2022-00817, F.H.S.) and the DAAD/ANID fellowship (57451854/62180003, J.C.). F.H.S. is supported by the Knut and Alice Wallenberg Foundation via the Wallenberg Centre for Molecular and Translational Medicine at the University of Gothenburg. Z.W. and C.Y. are investigators of SUSTech Institute for Biological Electron Microscopy.

## Author contributions

Z.W., C.A., F.H.S., and G.J. conceived the study. Z.W., C.A., and F.H.S. supervised the project. G.J., Y.L., J.C., and B.M.C. designed and performed experiments. L.L., J.D., C.Y., F.N., and Z.W. analyzed the data. Z.W., C.A., and F.H.S. wrote the manuscript with inputs from other authors.

## Competing Interests Statement

All authors declare that they have no competing interests.

## Data availability

The structure factors and atomic model of the liprin-α2_CC2N/RIM1_C2B complex have been deposited in the Protein Data Bank (PDB) with accession code 8Z22.

## Methods

### Constructs

Human liprin-α2 (GenBank: AF034799.1) CC12 truncation with an N-terminal His_6_-SUMO tag was generated in our previous study ^23^. The CC2 (residues 259 – 542), CC2N (residues 300 – 404), and CC2C (residues 405 – 542) regions were subcloned into a modified pET32a vector with N-terminal thioredoxin (Trx)-His_6_-tag and an HRV 3C protease cutting site. Plasmids encoding rat ELKS1 (Genbank: NM_170788.2), RIM1 (specifically the RIM1α isoform, Genbank: XM_017596673.1), and RBP2 (GenBank: XM_017598284.1) were kind gifts from Prof. Mingjie Zhang. For crystallization, RIM1_C2B (residues 1166 – 1334) was subcloned into a modified pET28a vector with an N-terminal His_6_-SUMO tag. Full-length RIM1 was inserted into the pCAG vector with an N-terminal FLAG tag. The full-length ELKS1 was first cloned into a modified pETL7 vector with an N-terminal followed by a TEV-protease cutting site. Subsequently, His_6_-MBP-GFP tagged ELKS1 was subcloned into the pCAG vector ^61^. All point mutations in these constructs were created using a site-directed mutagenesis kit. Lentiviral rescue constructs were generated by subcloning PCR-amplified wildtype and mutant liprin-α2 cDNA, using Gibson assembly, to a lentiviral vector containing the ubiquitin promoter. All constructs were verified by DNA sequencing. All plasmids used in this study were summarized in Table S2.

### Protein expression and purification

Trx or SUMO-tagged proteins were expressed in *Escherichia coli* BL21(DE3) cells. Transfected cells were cultured in LB medium at 37°C with agitation at 200 rpm to reach an OD600 of ∼0.8. After cooling to 16°C, protein expression was induced with 500 μM IPTG and continued with overnight shaking at 16°C and 200 rpm. Harvested cell pellets were lysed via high-pressure homogenization in a binding buffer (50 mM Tris pH 8.0, 500 mM NaCl, 5 mM imidazole) supplemented with 1 mM PMSF. The tagged proteins were purified using Ni^2+^-NTA affinity chromatography with an elution buffer (50 mM Tris pH 8.0, 500 mM NaCl, and 250 mM imidazole). The eluted proteins were further purified by size exclusion chromatography performed on a Superdex-200pg column (GE Healthcare) pre-equilibrated in TBS buffer (20 mM Tris pH 8.0, 100 mM NaCl, 1 mM EDTA, and 1 mM DTT). To prepare the RIM and liprin-α2 fragments for crystallization, the affinity tag was removed using HRV-3C or SUMO proteases at 4°C overnight, followed by a second round of size exclusion chromatography on a Superdex-75pg column (GE Healthcare) pre-equilibrated with TBS. Purified proteins were concentrated using Amicon Ultra centrifugal filters (Millipore) to ∼10 mg/mL, aliquoted and stored at -80°C after flash-freezing in liquid nitrogen. For fluorescence labeling, the Superdex-200pg column was equilibrated in a buffer containing 20 mM HEPES pH 8.0, 100 mM NaCl, 1 mM EDTA, and 1 mM DTT. The RBP2_(SH3)_3_ fusion protein was prepared as previously reported ^30^.

Full-length RIM1 and ELKS1 were expressed in HEK293F suspension cells (ThermoFisher Scientific), cultured in Freestyle 293 medium (OPM-293 CD05 Medium) at 37 ℃ supplied with 5% CO_2_ and 80% humidity. When cell density reached 2.0 × 10^6^ cells/mL, cells were transiently transfected using expression plasmids and polyethylenimine (PEI) (Yeasen Biotechnology). For transfection, ∼0.5 mg plasmids were pre-mixed with 1 mg PEIs in 50 mL fresh medium for 15 min, and then the mixture was added to 500 mL of cell culture. After a 72-hour culture, cells were collected at 4 ℃ by centrifugation at 1000 g for 20 min. The pellets were lysed in a buffer containing 50 mM Tris pH 7.5, 500 mM NaCl, 1 mM EDTA, 0.5% Triton X-100, and a protease inhibitor cocktail. Protein purification was performed using anti-FLAG affinity chromatography, with an elution buffer containing 100-500 μg of FLAG peptide (DYKDDDDK). Subsequent purification steps involved size-exclusion chromatography on a Superdex 6 Increase column (GE Healthcare) using TBS with varying NaCl concentrations according to the biochemical properties of individual proteins. The protein quality was further checked using an HT7700 transmission electron microscope (HITACHI) with 100 kV voltage.

### Co-immunoprecipitation (Co-IP) assay

Transfected HEK293T cells were lysed in ice-cold lysis buffer containing 50 mM Tris pH 7.5, 150 mM NaCl, 5% glycerol, 1% Triton X-100, 1 mM phenylmethylsulfonyl fluoride, 1% protease inhibitor cocktail (TargetMol, C001) for 0.5 h on ice and followed by centrifugation at 12,000 g for 15 min at 4°C. The supernatant fraction was then incubated with anti-GFP conjugated agarose beads (Ktsm-life, ktsm1301) for 60 min at 4°C. The beads were washed with the cell lysis buffer twice and resuspended with 20 μL SDS-PAGE loading buffer. The prepared samples were separated by 10% SDS-PAGE and transferred to polyvinylidene difluoride membranes (Millipore, IPVH00010). The membranes were sequentially blocked with 5% skim milk in buffer containing 50 mM Tris-HCl pH 7.4, 150 mM NaCl, and 0.1% Tween 20, immunoblotted with anti-GFP mouse monoclonal antibody (Transgen, HT801-01, dilution 1:3000) or anti-DYKDDDDK(anti-FLAG) mouse monoclonal antibody (Transgen, HT201-01, dilution 1:3000), probed with horseradish-peroxidase conjugated secondary antibodies (Cell Signaling, 7076s, dilution 1:10000) at room temperature and finally developed with a chemiluminescent substrate (BioRad, 107-5061). Protein bands were visualized on the Tanon-6011C Chemiluminescent Imaging System (Tanon Science and Technology).

### Isothermal titration calorimetry (ITC) assay

To quantitatively analyze protein-protein interaction, ITC experiments were conducted using a MicroCal PEAQ-ITC calorimeter (Malvern Panalytical). All proteins were prepared in an identical reaction buffer containing 20 mM Tris pH 8.0, 100 mM NaCl, and 1 mM EDTA. The protein concentration in the syringe was 400 μΜ for titrating into the reaction cell, where the concentration of target proteins was typically 40 μΜ. Experiments were carried out at a controlled temperature of 25°C. Each titration involved injecting 3 μL of the syringe solution into the cell, followed by a 150-second equilibration period between injections. A titration curve contained a total of 13 titration points. The resulting data were analyzed using the MicroCal PEAQ-ITC Analysis software, applying a one-site binding model to determine the dissociation constant (*K*_d_).

### Analytical size exclusion chromatography (aSEC)

Analytical gel filtration chromatography was performed using an ÄKTA Pure system (GE Healthcare). The protein samples were loaded into a Superdex 200 Increase 10/300 GL column (GE Healthcare) pre-equilibrated with a buffer comprising 20 mM Tris-HCl pH 7.5, 100 mM NaCl, 1 mM EDTA, and 1 mM DTT.

### SEC coupled with multi-angle light scattering (SEC-MALS) assay

The SEC-MALS assay was conducted using a platform composed of a multi-angle light scattering (MALS) detector (miniDawn, Wyatt), a differential refractive index (dRI) detector (Optilab, Wyatt) and a liquid chromatography (LC) system (AKTA pure, GE Healthcare). In each assay, a 100 μl sample (individual proteins or complexes) was injected into a Superdex 200 Increase 10/300 GL column (GE Healthcare) pre-equilibrated with TBS. Data were analyzed using ASTRA6 (Wyatt).

### Protein crystallization and structure determination

Crystals of the liprin-α2_CC2N/RIM1_C2B complex were grown using the sitting drop vapor-diffusion method. Protein samples of liprin-α2_CC2N and RIM1_C2B were mixed in a 1:1 ratio and concentrated to 21 mg/mL. This concentrated mixture (1 μL) was combined with an equal volume of reservoir buffer containing 28% v/v 2-propanol, 0.1 M BIS-TRIS pH 6.5, and 3% v/v polyethylene glycerol 200 for crystallization tray setup. The crystallization was conducted at 16°C, and the resulting crystals were cryoprotected with 30% (v/v) glycerol. X-ray diffraction data were collected at the BL19U1 beamline of the Shanghai Synchrotron Radiation Facility (SSRF). Diffraction data were processed using HKL2000 software ^62^. The complex structure was solved by molecular replacement in PHASER ^63^ using the RIM1_C2B apo structure (PDB ID: 2Q3X) as the search model. Model building, adjustment, and refinement were carried out iteratively using COOT ^64^ and PHENIX ^65^. The final models were validated by MolProbity ^66^ and statistics were summarized in Table S1. All structure figures presented in the paper were prepared using PyMOL (https://www.pymol.org/).

### Fluorophore labeling of proteins

Fluorescent labeling dyes, including Cy3/Cy5/iFluor 488 NHS esters (ThermoFisher) and iFluor 405 NHS ester (AAT Bioquest), were dissolved in DMSO at a stock concentration of 5 mM and stored at -20°C. Prior to labeling, proteins were concentrated at 5 mg/mL in a HEPES buffer at pH 7.5 to ensure specific N-terminal labeling. Labeling was performed by mixing the proteins with the corresponding fluorophores at a 1:1 molecular ratio and incubating at room temperature for 1 hour. The reaction was quenched by adding a 200 mM Tris buffer. Unincorporated fluorescence was removed using a pre-equilibrated HiTrap desalting column (GE Healthcare) with the corresponding TBS buffer. Fluorescence labeling efficiency was assessed using a Nanodrop-2000 spectrophotometer (ThermoFisher). The labeled proteins were frozen and stored at -80°C. For imaging, a sparse labeling approach was used, where the fluorescence-labeled proteins were mixed with an excess of corresponding unlabeled proteins in the same buffer, achieving a final molecular ratio of 1:100.

### Phase separation assays

#### Sample preparation and imaging

Prior to imaging experiments, all proteins used for imaging were centrifuged at 20,000 g for 10 min at 4°C to remove any potential aggregates or precipitates. Protein concentrations and buffer conditions were specified in the corresponding figures or their legends. For imaging, samples were applied to the wells of 384-well glass bottom plates (P384-1.5H-N, Cellvis). Confocal images were captured using an A1R confocal microscope (Nikon) equipped with a 100X/NA oil objective lens. The fluorescence intensities of images were analyzed using ImageJ/Fiji software. For the phase separation assay involving RIM1, RBP2_(SH3)_3_, and liprin-α2_CC12, RIM1α and RBP2_(SH3)_3_ were mixed at the desired concentration in a 200 μL microcentrifuge tube, then applied to the 384-well plates. After allowing the condensates to settle for 10 minutes, the CC12 fragment or its mutant was introduced into the well. Following an additional 5-minute incubation, images were captured. For the phase separation assay involving both RIM1 and ELKS1 condensates, the MBP tag on the ELKS1 fusion protein was removed using TEV protease to produce GFP-ELKS1 condensates. After RIM1/RBP2_(SH3)_3_ condensates had settled down for 10 min, ELKS1 condensates w/o liprin-α2_CC12 were added to the 384-well plate and mixed with RIM1α/RBP2_(SH3)_3_ condensates. The mixtures were then allowed to settle for an additional 20 min before image capture. To detect the distribution of NCav_CT in the two-phase system, NCav_CT was mixed with ELKS1 condensates and then loaded onto the pre-formed RIM1α/RBP2_(SH3)_3_ condensates.

#### Sedimentation assay

RIM1 was mixed with RBP2_(SH3)_3_/liprin-α2_CC12 at the specified concentration and incubated for 5 min. The mixture was centrifuged at 1,400 rpm for 5 min at room temperature. Following centrifugation, the supernatant was promptly isolated by pipette thoroughly, and the pellet was re-suspended with 20 μL of dilution buffer. Both supernatant and pellet samples on the SDS-PAGE gel were visualized using Coomassie blue R250 staining. The intensity of bands of interest was quantified using ImageJ/Fiji software.

#### Fluorescence recovery after photo-bleaching (FRAP) assay

In each FRAP experiment, ten regions of interest (ROIs) were selected. Laser beams at 561nm with 100% power were precisely applied to target the Cy3 fluorophore for photobleaching. Pre-bleach and post-bleach images were acquired with no delay time interval. Subsequent time-lapse images were captured at 20-second intervals for a duration of 20 min to record fluorescence intensity recovery. These experiments were conducted using a Nikon A1R confocal microscope equipped with a 100X/NA oil lens. Fluorescence recovery was measured with ImageJ/Fiji by calculating the intensity at each time point. Data were processed by correcting for background fluorescence and normalizing the pre-bleaching intensity to 100% and the bleaching point intensity to 0%.

### Cell culture experiments

#### Maintenance of human embryonic stem cells (hESCs)

hESCs of line WA09/H9 (RRID: CVCL_9773) were obtained from WiCell and maintained/cultured on Matrigel-coated (Corning #15505739) dishes using mTeSR Plus (StemCell Technologies #100-0276), which was changed every other day. hESC cells were kept in an incubator supplied with 5% CO2 at 37°C. All procedures followed The Robert Koch Institute guidelines for human ESC work.

#### Maintenance of human embrynic kidney (HEK) cells

HEK (HEK293T/17, ATCC CRL-11268) cells were used to produce all lentiviruses for this study. Cells were maintained in an incubator supplied with 5% CO2 at 37°C, using DMEM-Glutamax medium (Gibco #31966047) supplemented with 10% fetal bovine serum (FBS; Sigma #F7524). The medium was changed every other day, and cells were split after reaching near 70% confluence using Trypsin-EDTA (Gibco #15400054).

#### Production of lentiviruses for neuronal differentiation, optogenetic control, and sparse visualization

Lentiviruses were produced in HEK cells, as described elsewhere ^67^. Briefly, two hours before transfection, at near 60% confluence, the medium was changed, and then HEK cells co-transfected using Linear Polyethylenimine 25,000 (PEI, Sigma Cat# 23966) with the following plasmids: pREV (3.9 μg), pRRE (8.1 μg), pVSVG (6 μg), and with the corresponding vector DNA using 12 μg per 75 cm^2^ cell culture area. The following lentiviral vector DNA was used to produce lentiviruses for differentiation, optogenetic control, and sparse visualization: FU-M2rtTA, Tet-O-*Ngn2*-puromycin, Channel-rhodopsin oChiEF fused to tdTomato (termed here ChR-tdTomato), and soluble GFP ^26^. Two hours after transfection, the medium was changed again with fresh DMEM media, and lentiviruses were harvested from the medium 40 hours later. Specifically, the medium was first centrifuged at 1,500 g for 10 min at 4°C to eliminate dead cells and debris, and then lentiviral particles were pelleted by high-speed centrifugation (60,000 g for 1.5 h), resuspended in MEM (Gibco #51200046) with 10 mM HEPES (100 μl per 30 ml of medium), aliquoted, and snap-frozen in liquid nitrogen.

#### Production of lentiviruses for neuronal infection (rescue constructs)

All rescue experiments were performed using fresh (non-concentrated) lentiviruses, generated using the same protocol described above but with the following modifications. First, two hours after transfection, the medium was replaced with Neurobasal supplemented with 2% B27 (Gibco #17504044), GlutaMAX (Gibco #35050061), and 10 mM HEPES (Gibco #15630080). Second, lentiviral particles were harvested from the medium 40 h after transfection with a low-speed (1,500 g for 10 min at 4°C) centrifugation to pellet dead cells and debris. The supernatant was then aliquoted and frozen at -80°C.

#### Generation of induced neurons

Induced glutamatergic neurons were generated from control (Ctrl1) and liprin-α1 to α4 mutant (qKO1) ESC clones, as described in detail previously ^68^. In brief, for each neuronal experiment, 250K hESCs were detached with Accutase (Gibco), plated on matrigel-coated wells in mTeSR Plus containing Rho kinase inhibitor (Y27632, Axon Medchem #1683), and transduced with concentrated lentiviruses FU-M2rtTA and Tet-O-*Ngn2*-puromycin, generated as described in the previous section. A day later (defined as DIV0), the medium was changed to N2 medium [DMEM/F12 (Gibco #11330032), 1% N2 supplement (Gibco 17502048) 1% non-essential amino acids (Gibco #11140050), laminin (200 ng/ml, Thermo Fisher #23017015), BDNF (10 ng/ml, Peprotech #450-02) and NT-3 (10 ng/ml, Peprotech #450-03) supplemented with Doxycycline (2 μg/ml, Alfa Aesar)] to induce expression of *Ngn2* and the puromycin resistance cassette. On DIV1, puromycin (1 mg/ml) was added to the medium, and after 48 hours, selection cells were detached with Accutase (Gibco #A1110501) and re-plated on Matrigel-coated coverslips along with mouse glia (typically at a density of 150,000 iGluts per 24-well plate) in B27 medium [Neurobasal-A (Gibco #12349015) supplemented with B27 (Gibco #17504044), GlutaMAX (Gibco #35050061) laminin, BDNF, and NT-3]. From this time point on until DIV10, the medium was replaced every second day, and cytosine arabinoside (ara-C; Sigma #C6645) was added to a final concentration of 2 μM to prevent glia overgrowth. Rescue lentiviral constructs (e.g., to express either WT or mutant liprin-α2 constructs that prevent interaction with RIM) were added to the medium on day 4. From DIV10, neuronal growth medium [Neurobasal-A supplemented with B27, GlutaMAX, and 5% FBS (Hyclone #SH30071.03HI)] was washed in and used for partial medium replacements every 3-4 days until analysis, typically at around 6 weeks in culture.

#### Induced neurons for measuring evoked transmission

To measure the impact of the slow calcium buffer EGTA on evoked synaptic transmission, the protocol for the generation of induced glutamatergic neurons was slightly modified, as it required the majority (80%) of cells to express ChR and a minority (20%) to express GFP for visualization for electrophysiological recordings. For this, cells from either control or qKO clones were further separated into two groups. In group #1, cells were infected with pFU-M2rtTA, pTet-O-*Ngn2*-puromycin, and lentiviruses expressing ChR-tdTomato ^46,68^. In group #2, cells were infected with pFU-M2rtTA, pTet-O-*Ngn2*-puromycin, and lentiviruses to express soluble GFP. Four days later, cells from groups #1 and #2 were washed three times with PBS to remove any lentivirus trace attached to cell membranes, detached, mixed at a ratio of 80/20 (80% with ChR and 20% with GFP), re-seeded on Matrigel-coated coverslips along with mouse glia, and cultured as described above.

#### Isolation of mouse glial cells

For induced glutamatergic neurons to form mature functional synapses, we grew induced neurons on a monolayer of primary mouse glial cells. Isolation and culture of primary mouse glial cells were performed essentially as described ^68^. In short, E21-P1 mouse cortices from wildtype C57BL6 mice were dissected and triturated with fire-polished pipettes and filtered through a cell strainer. Cells from two cortices were plated onto T75 flasks pre-coated with poly-L-lysine (5 mg/mL, Sigma #P1274) in DMEM supplemented with 10% FBS (Sigma). Upon reaching confluence, cells were dissociated by trypsinization and re-seeded twice to remove potential trace amounts of mouse neurons before the glia cell cultures were used for co-culture with induced neurons.

#### Immunocytochemistry

Cultured neurons were fixed for 15 min at room temperature with a solution containing 4% Paraformaldehyde and 4% Sucrose in PBS, pH 7.4, then washed three times with PBS (each washing step was separated by 10 mins) and permeabilized with 0.1% TritonX-100 in PBS for 10 min at room temperature. Then, cultures were blocked for 1 hour (2% NGS, 1% BSA, 0,01% NaN_3_ in PBS), and incubated in primary antibodies diluted in the blocking buffer overnight at 4°C in a humidity chamber. The next day, neurons were washed (3×) with PBS and incubated with fluorescently labeled secondary antibodies (Jackson ImmunoResearch) for 1 hour at room temperature. After this, neurons were washed 3 times in PBS, followed by a wash in ddH2O and mounting in microscope slides using ProLong Gold mounting medium (Thermo Fisher Scientific). For PSD95 staining, a modified version of the protocol described above was utilized. Specifically, neurons were fixed in ice-cold Methanol fixing solution (90% methanol and 10% MES buffer: 100 mM MES pH 6.9, 1 mM EGTA, and 1 mM MgCl_2_) at room temperature for 5 min, washed 3 times in PBS, and then incubated in blocking/permeabilizing solution (2% normal goat serum, 1% BSA 0.01% NaN_3_, and 0.1% Triton X-100 in PBS) for 30 min, before proceeding with staining. The following primary antibodies and dilutions were used: MAP2 (Encor, 1:1000), pan-Synapsin (Proteogenix, 1:1000), PSD-95 (NeuroMab, 1:100 and Addgene 1:100 for SIM experiments), RIM1/2 (SySy, 1:200), RIMBP-2 (SySy, 1:200), Synaptophysin-1 (SySy, 1:200), Piccolo (SySy, 1:200), Cav2.1 (SySy, 1:200). See Table S3 for details on the source of each antibody.

### Confocal imaging

For measurements of synapse density and gross estimations of active zone integrity, synapse-rich areas were identified by extensive synapsin signals in close proximity to postsynaptic MAP2 profiles and were imaged using a confocal microscope (Leica, Germany) controlled by LAS X software (Leica, Germany). Specimens were sampled in frame mode at 1024×1024 pixels/frame resolution. 10 optical sections along the z-axis were taken for each sample and then compiled into a single maximal projection image for analysis. All the acquisition parameters were kept constant between conditions and experiments.

### Super-resolution STED imaging

STED imaging was conducted similarly as described previously ^69^. Briefly, induced neurons were cultured on glass coverslips, and immunocytochemistry was performed as described above, except that five PBS washing steps were done after each antibody incubation and Alexa 488 anti-guinea pig (Thermo), STAR 580 nanobody anti-mouse (Abberior) and STAR 635P anti-rabbit were used as secondary antibodies. Image acquisition was conducted in a Leica SP8 Confocal/STED 3× microscope with an oil immersion 100×1.44 numerical aperture objective and gated detectors (2-6 ns range for all 3 signals). Images were acquired from synapse-rich areas of 33.2 μm^2^ sampled at ∼16 nm per pixel. The signal from the 488 antibodies was acquired in confocal mode, and signals from the 580 and the 635 antibodies were acquired in STED mode sequentially (to avoid bleed trough) and using the same STED laser line (775 nm to maximize alignment). Line accumulation (4-8×) and frame averaging (3×) were applied. Images were acquired blindly to the genotype of the samples and identical settings were used for all the samples within an experiment/batch.

### Western blot

Protein samples were extracted from cultured neurons at DIV30-45, lysed in RIPA buffer (50 mM Tris pH 8.0, 150 mM NaCl, 0.1% sodium dodecyl sulfate, 0.5% sodium deoxycholate, and 1% Triton X-100) supplemented with PMSF (Thermo Fischer #36978) and Complete proteinase inhibitor cocktail (Merch #11873580001) for 20 min. Lysates were centrifuged at 20,000 g for 10 min at 4°C, and supernatants containing solubilized proteins were collected. Protein samples were separated by SDS-PAGE in pre-cast TGX gels (Biorad). Transfer to a nitrocellulose membrane (Amersham) was performed in Towbin transfer buffer (25 mM Tris, 0.2 M glycine, and 20% methanol). Membranes were blocked with 5% non-fat milk (Aplichem) for 1 hour and primary antibodies were incubated overnight at 4°C. After washing the membranes three times with TBS-T (20 mM Tris pH 7.5, 137 mM NaCl, and 0.05% Tween-20), secondary antibodies were incubated in 1:1 TBS-T Odyssey Blocking (LI-COR # 927-50000) for 1 hour. Membranes were imaged using an Odyssey CLx system (LI-COR), and bands were quantified by densitometry using Image Studio 5.2 software (LI-COR). The following primary antibodies (see Table S3 for details) were used: Tuj1 (BioLegend, 1:5000) and GFP (Thermo Fisher Scientific, 1:1000).

### Patch clamp electrophysiology

#### General

All electrophysiological recordings were done using an RC-27 chamber (Sutter Instruments) mounted under a BX51 upright microscope (Olympus). The microscope was equipped with DIC and fluorescent capabilities, and with a TTL-driven LED for optogenetic activation with millisecond precision. All recordings were done at 26 ± 1°C using a dual TC344B temperature control system (Sutter Instruments), with neurons continuously perfused with oxygenated (95% O2 and 5% CO2) ASCF containing 125 mM NaCl, 2.5 mM KCl, 0.1 mM MgCl_2_, 4 mM CaCl_2_, 25 mM glucose, 1.25 mM NaH_2_PO_4_, 0.4 mM ascorbic acid, 3 mM myo-inositol, 2 mM Na-pyruvate, and 25 mM NaHCO3, pH 7.4. Neurons were approached and patched under DIC, using 3.0 ± 0.5 MegaOhm pipettes (WPI), pulled with a PC10 puller (Narishige, Japan). In all experiments, pipettes were loaded with a voltage clamp internal solution containing 125 mM Cs-gluconate, 20 mM KCl, 4 mM MgATP, 10 mM Na-phosphocreatine, 0.3 mM GTP, 0.5 mM EGTA, 2 mM QX314 (Hello Bio, #HB1030), and 10 mM HEPES, pH 7.2. Electrical signals were recorded using a Multiclamp 700B amplifier (Axon Instruments) controlled by Clampex 10.1 and Digidata 1440 digitizer (Molecular Devices).

#### Spontaneous synaptic current recordings

We performed whole-cell voltage-clamp recordings at ∼70 mV holding potentials. Spontaneous excitatory currents were detected as downward deflections from the baseline.

#### Evoked currents

Evoked excitatory currents were triggered by 5 ms pulses of blue (488 nm) light generated via a CoolLED illumination system (pE-300) controlled by a TTL pulse, and recorded from GFP+/ChR-neurons (see above) in voltage clamp at ∼70 mV holding potentials.

#### Sucrose responses

Cells were maintained at ∼70 mV holding potentials (voltage clamp configuration) and stimulated with 0.5 M sucrose solution for 5 seconds. Sucrose solution was delivered in the vicinity of recorded cells (20-30 μm away) using a low resistance glass pipette (1.5 MegaOhms) connected to a custom pressure device (5 psi).

### Data analysis and statistics

Confocal images were handled and analyzed using LASX (Leica) or ImageJ/Fiji (v. 2.3.0/1.53f). In experiments aimed to measure the recruitment of RIM1, Piccolo, and RBP2 to presynaptic boutons in qKO neurons, the signal intensity of corresponding active zone markers was measured only inside ROIs defined by Synaptophysin. For STED image analysis, individual “side view” synapses were manually selected, and intensity profiles were obtained by drawing a rectangle of 1200×200 nm centered in and perpendicular to the PSD-95 elongated signal using an ImageJ custom macro. Intensity profiles were recorded for all 3 signals, and the right alignment/orientation of the profiles was performed in R Studio. Intensity traces were obtained by averaging individual traces over the raw data values. Representative images in figures were linearly adjusted using bright and contrast identically across samples. Immunoblot images were handled and analyzed with Image Studio v. 5.2 (LI-COR). Analysis of voltage-and current-clamp recordings was done with Clampfit 10.1 or with custom-written macros in IgorPro 6.11. Electrophysiological/imaging experiments were done and analyzed with the experimenter blinded to the sample genotype/condition whenever possible. Summary data are shown as means ± standard errors of the mean (SEM) or means ± standard deviation (SD) as indicated in the figure legends. Statistical analysis was performed using Prism 9 (GraphPad Software). Datasets were first analyzed with the D’Agostino Pearson test to determine if the data had a normal (Gaussian) distribution. For between-group comparisons, unpaired two-tailed t-tests were used if data distribution was normal, or two-tailed Mann-Whitney tests for non-Gaussian datasets. For multiple-group comparisons, statistical significance was determined by ANOVA with Tukey’s or Holm-Šídák’s corrections for multiple comparisons, or Kruskal-Wallis (KW) followed by Dunn’s post hoc test for non-Gaussian datasets. ns, not significant; ****, p < 0.0001; ***, p < 0.001; **, p < 0.01; *, p < 0.05.

## Supplementary Figures

**Figure S1.**
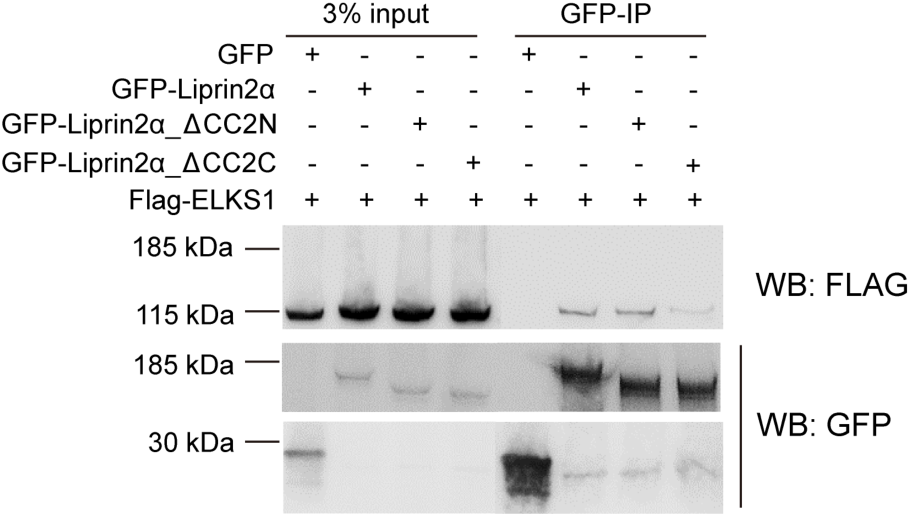
Co-immunoprecipitation analysis of the CC2-mediated binding of liprin-α2 to ELKS1.

**Figure S2.**
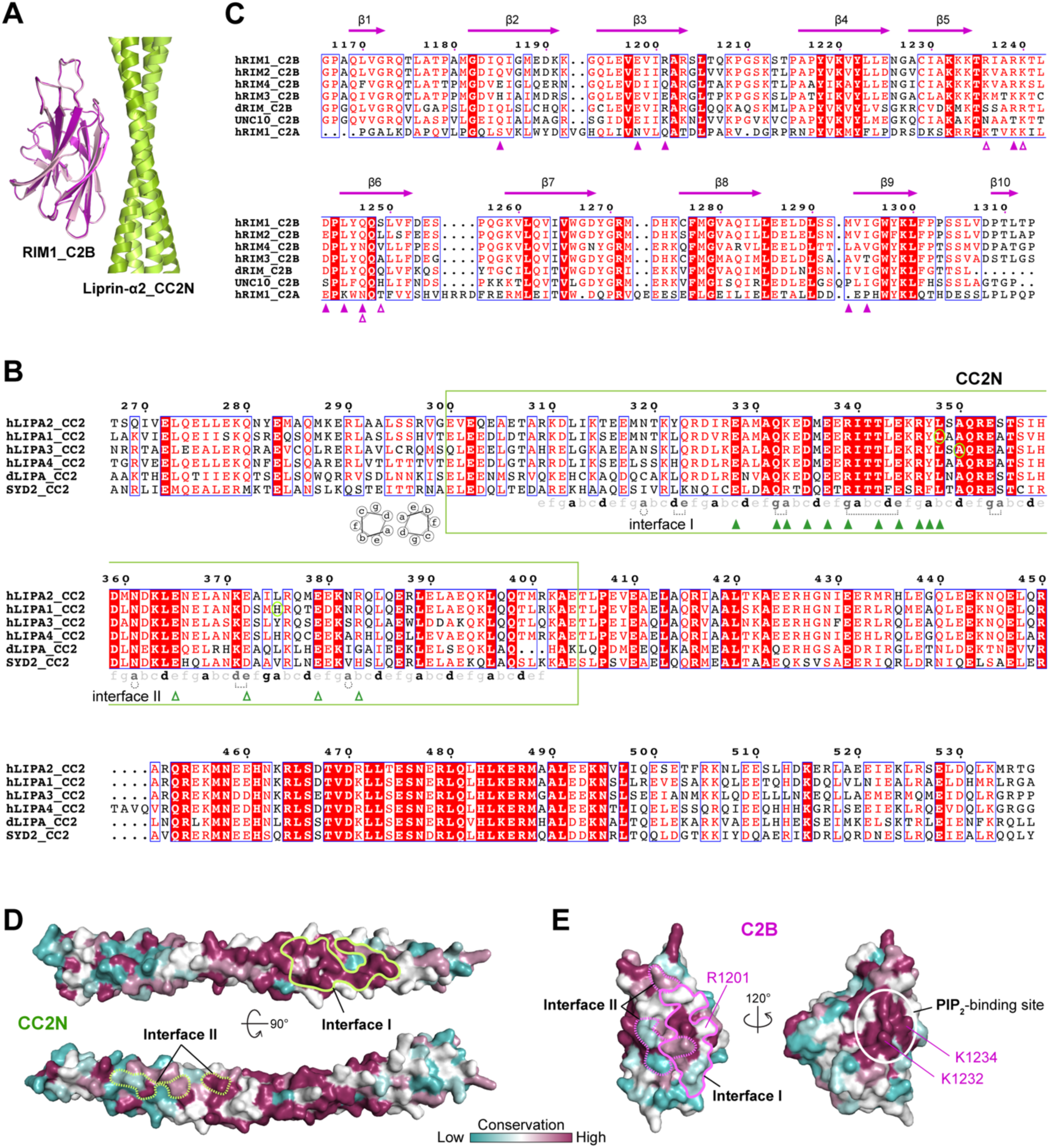
Structural and sequence analysis of the liprin-α2/RIM1 interaction. **A.** Structural comparison of the two C2B molecules (e.g., C2B and C2B’ in Fig. 2A) in the CC2N/C2B tetrameric complex. The corresponding interfaces I on CC2N are aligned well, indicating the symmetric binding of C2B to the dimeric CC2N coiled coil. **B.** Multisequence alignment of the CC2 segment from liprin-α family proteins. Species abbreviations: ’h’ for human, ’d’ for Drosophila, and SYD2 as the *C.elegans* liprin-α homolog. Heptad repeats of coiled-coil structures in the CC2N sequence are annotated, with heptad labels in black and dark grey denoting residues involved in coiled-coil formation through hydrophobic and polar interactions, respectively. Polar interactions are indicated by dashed lines, while residues involved in interfaces I and II are marked by solid and open purple triangle-ups. Mutation sites associated with neurodevelopmental disorders are encircled. **C.** Multisequence alignment of the C2B domains from RIM family proteins. The sequence of the C2A domain in human RIM1 was also included in the alignment for comparison. UNC10 is a RIM homolog in *C.elegans*. Residues involved in interfaces I and II are indicated by solid and open green triangle-ups, respectively. **D.** Surface conservation analysis of the CC2N coiled-coil structure. Conservation scores for each residue were calculated based on the alignment in panel **B**. **E.** Surface conservation analysis of the C2B structure. Conservation scores for each residue were calculated based on the alignment in panel **C**. The highly conserved PIP_2_-binding site is indicated by a circle.

**Figure S3.**
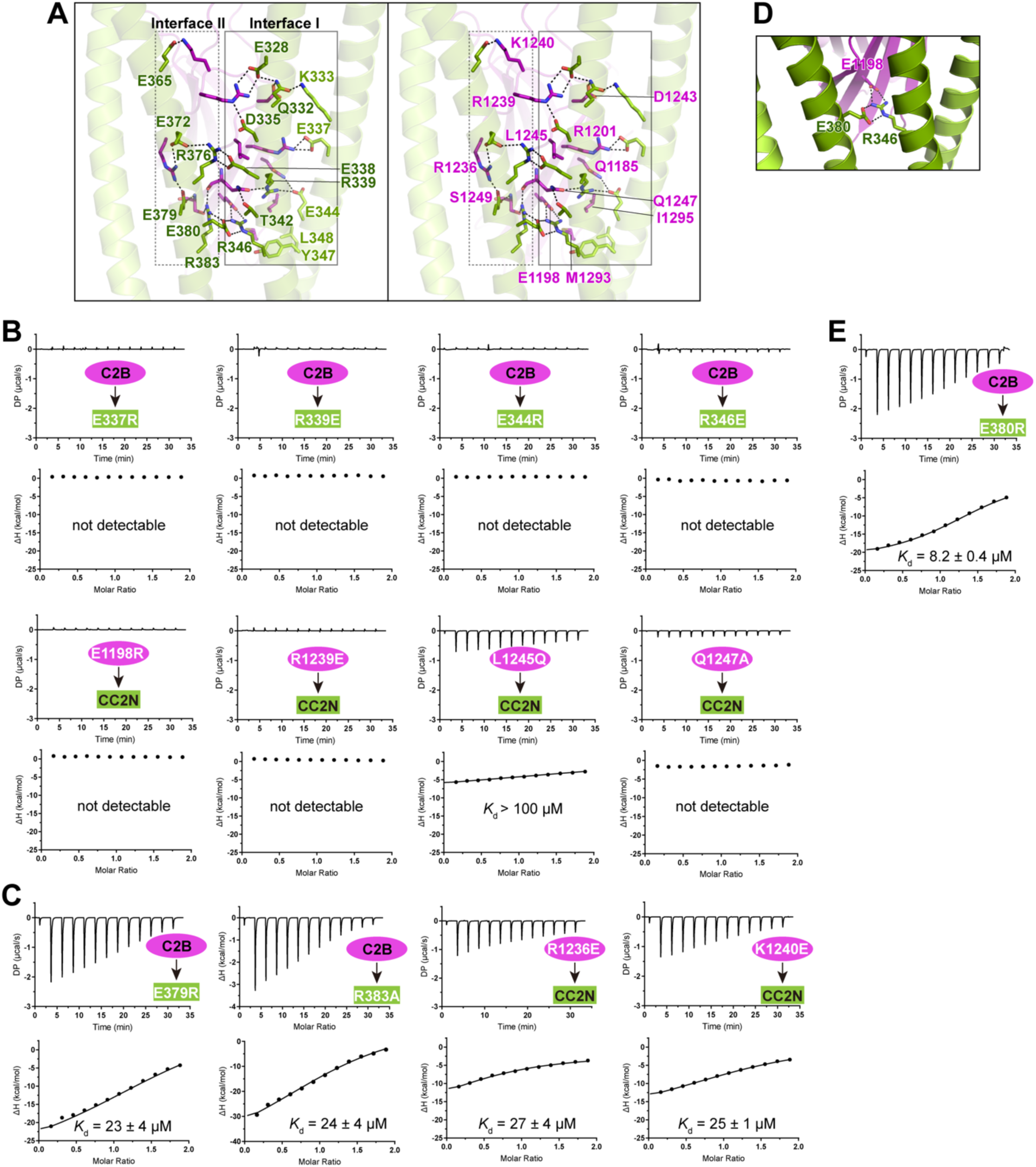
Structure-based mutagenesis and ITC-based analysis of the liprin-α2/RIM1 interaction. **A.** Stereo view showing the molecular details of the coupling between the two interfaces of CC2N and C2B. Key salt bridges and hydrogen bonds are indicated by dashed lines. **B.** ITC-based analyses of interface I mutation effects on the CC2N/C2B interaction. **C.** ITC-based analyses of interface II mutation effects on the CC2N/C2B interaction. **D.** Structural analysis of two interacting CC2N coiled coils at interface I. The salt bridge formed between R346^liprin-α2^ and E1198^RIM1^ is stabilized by E380^liprin-α2^. **E.** ITC-based analysis showing the mild disruptive effect of the E380R mutation in CC2N on the CC2N/C2B interaction.

**Figure S4.**
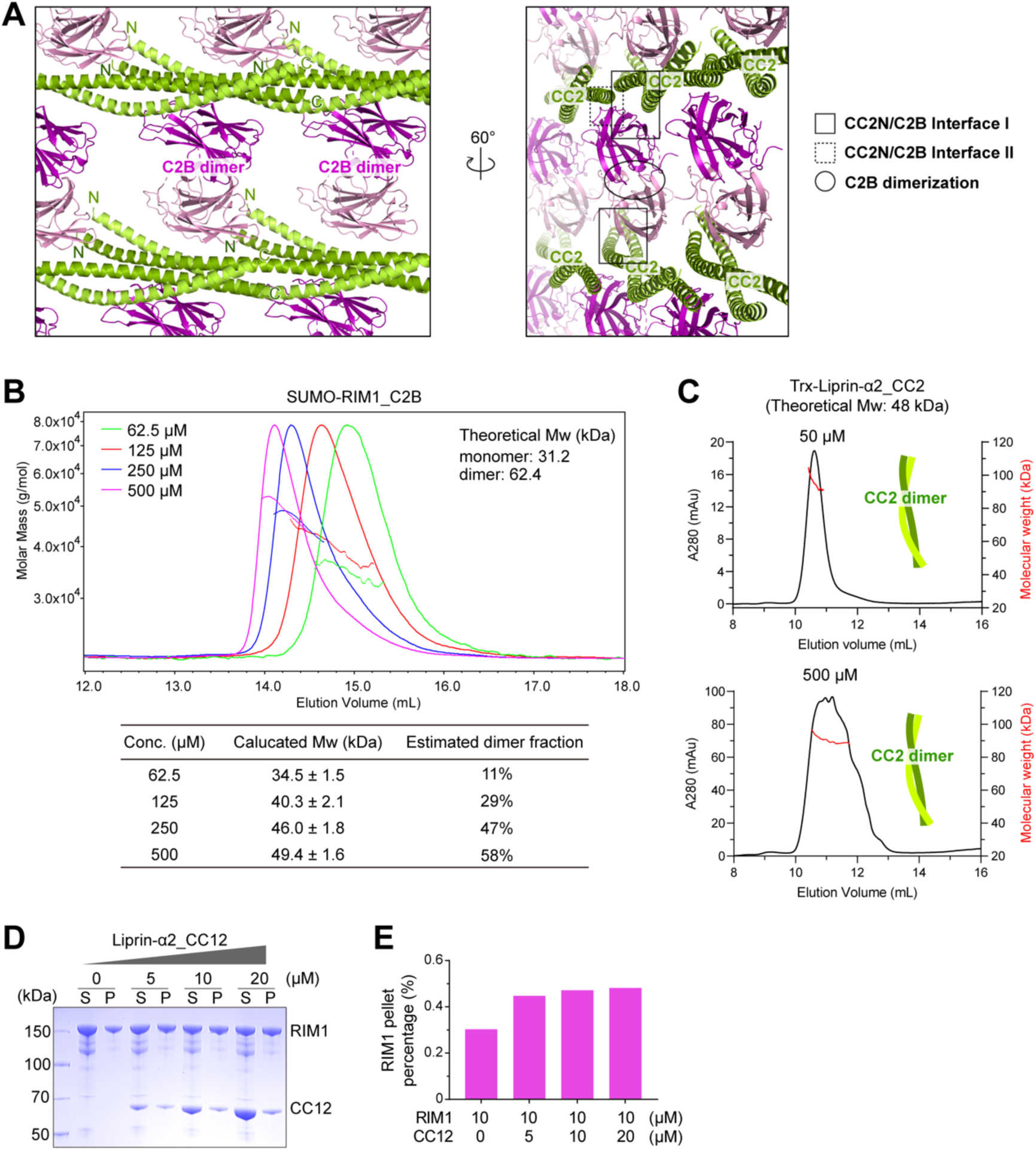
Analyses of multiple intermolecular interactions in the liprin-α2/RIM1 assembly. **A.** Crystal packing analysis of the liprin-α2_CC2N/RIM1_C2B complex. Different intermolecular interfaces are highlighted. **B.** Dimerization fraction analysis of RIM1_C2B in solution using aSEC coupled with MALS. **C.** Molecular weight analysis of liprin-α2_CC2 in solution using aSEC coupled with MALS. **D.** Sedimentation-based assay indicating the distribution of RIM1 full-length protein in the supernatant (S) and pellet (P) when mixed with increasing concentrations of liprin-α2_CC12. **E.** Quantification of the RIM1 content of pellet fraction of samples shown in panel **D**.

**Figure S5.**
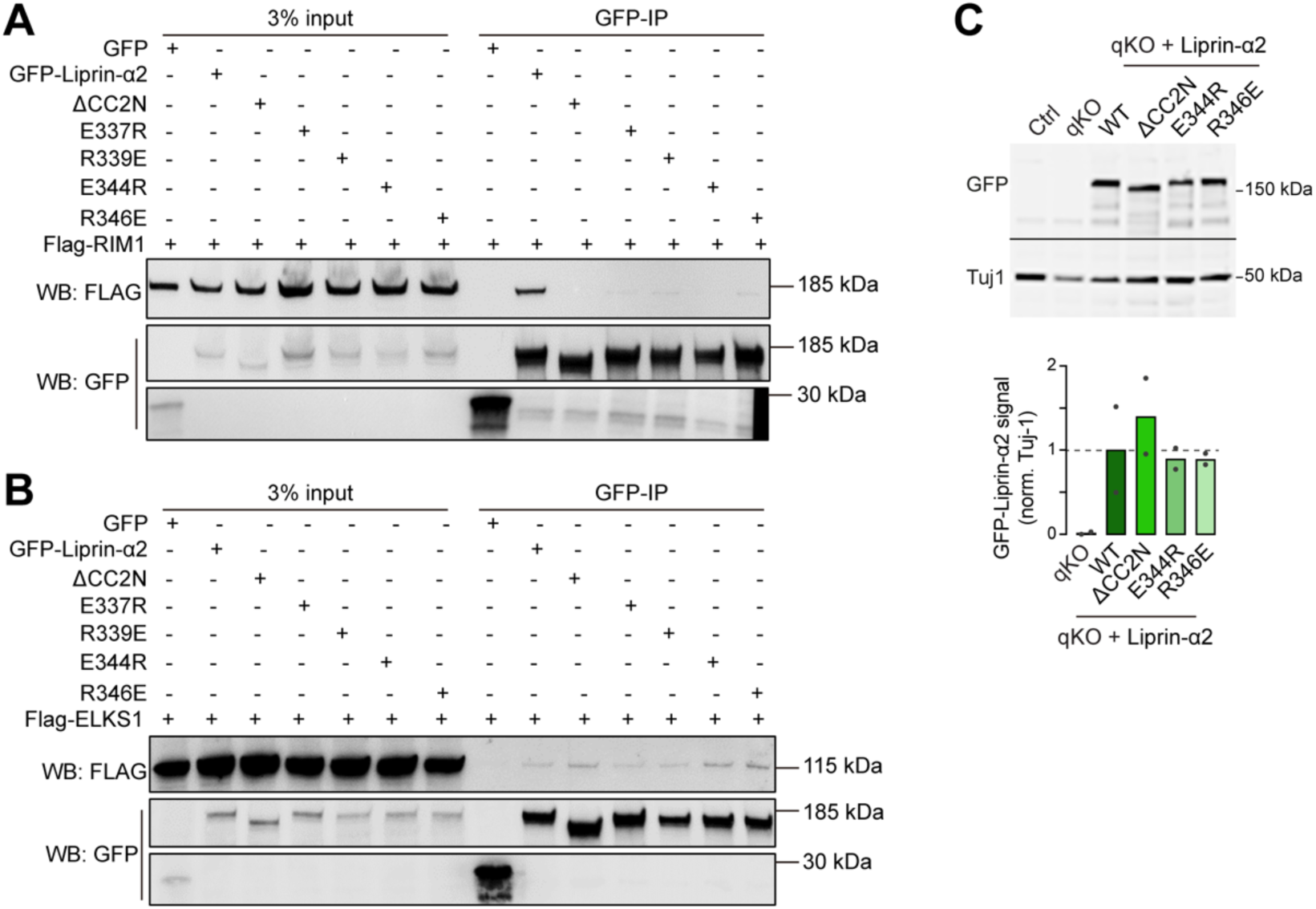
Characterization of liprin-α2 mutations for rescue assays in liprin-α qKO neurons. **A.** Co-immunoprecipitation analysis of the binding of liprin-α2 variants to RIM1. Consistent with our ITC-based analyses (Fig. S3B), the interface I mutations in liprin-α2 disrupt the liprin-α2/RIM1 interaction. Results are repeated by three independent batches of experiments. **B.** Co-immunoprecipitation analysis of the binding of liprin-α2 variants to ELKS1. The interface I mutations, especially E344R and R346E in liprin-α2, showed minimal interference with ELKS1 binding. Thus, these mutations were selected for the following rescue assays in liprin-α qKO neurons. **C.** Western blot analysis confirming comparable expression levels of liprin-α2 variants with lentivirus transfection in liprin-α qKO neurons.

**Figure S6.**
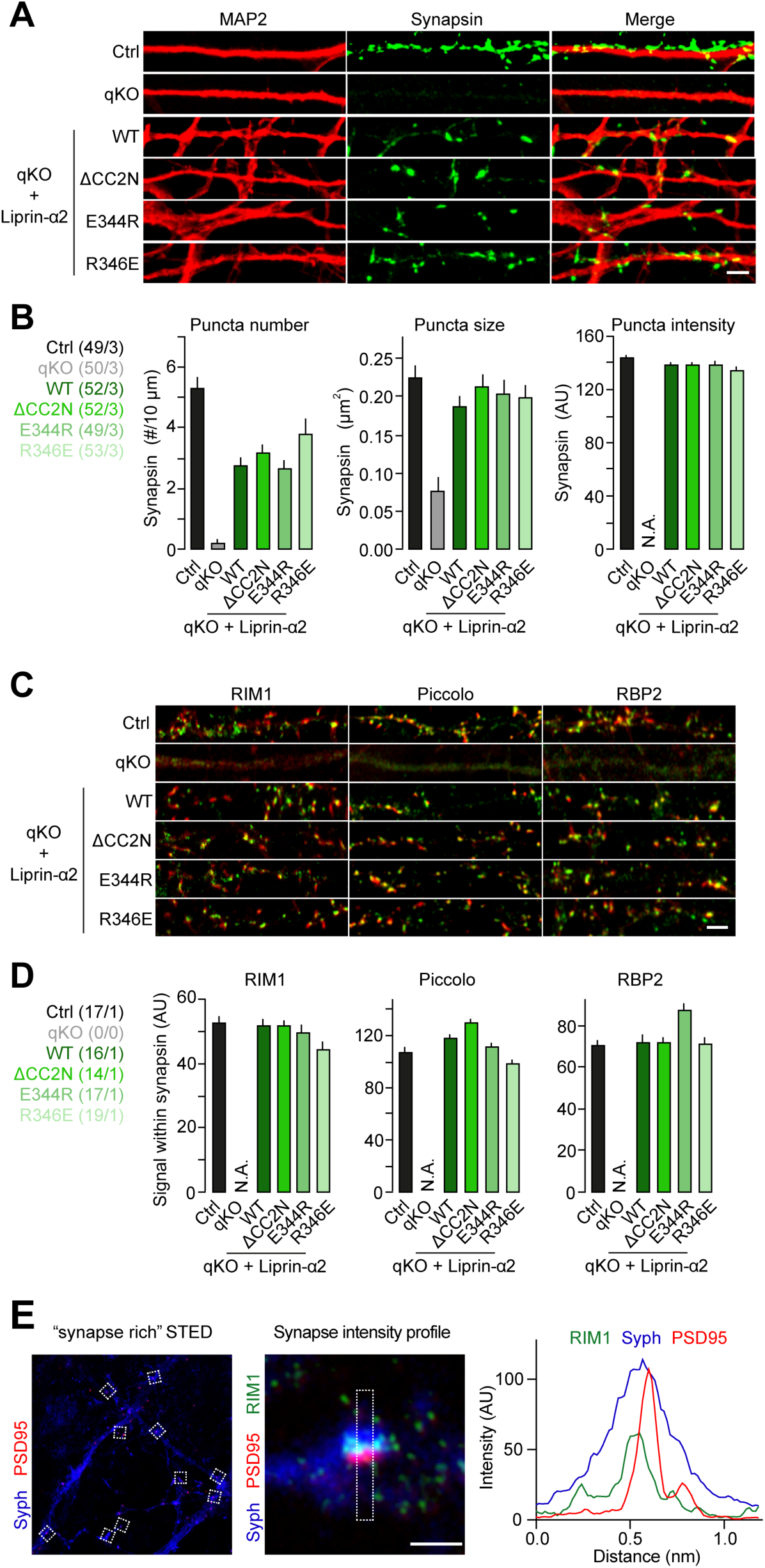
Morphological integrity of human synapses upon disruption of liprin-α/RIM complexes. **A.** Representative confocal micrographs showing presynaptic synapsin (green) puncta closely apposed to dendritic MAP (red) profiles under the indicated conditions. Scale bar: 5 μm. **B.** Quantification of synapsin puncta density (left), size (middle), and intensity (right). The data reflects the minimal impact of liprin-α/RIM complex perturbations on synapse morphological integrity. The number of cells/batches analyzed for each condition is indicated on the right. **C.** Confocal micrographs displaying the distribution of active zone proteins Piccolo (left), RIM (middle), and RBP2 (right) at presynaptic boutons under the indicated conditions. Scale bar: 5 μm. **D.** Quantification of fluorescence intensity from experiments presented in panel **C**. Signal intensity of active zone markers was measured only inside regions of interest (ROIs) defined by synapsin staining (see Methods for details). The number of ROIs/batches analyzed for each condition is indicated on the right. **E.** Microscopic analysis to identify synapses and measure the sub-synaptic distribution of presynaptic proteins using STED microscopy. Synapse-rich regions identified by Synapsin/PSD95 appositions were visualized at low magnification (left), followed by high magnification imaging (middle), with signal intensity quantified relative to the distance from postsynaptic PSD95 signals (right).

**Figure S7.**
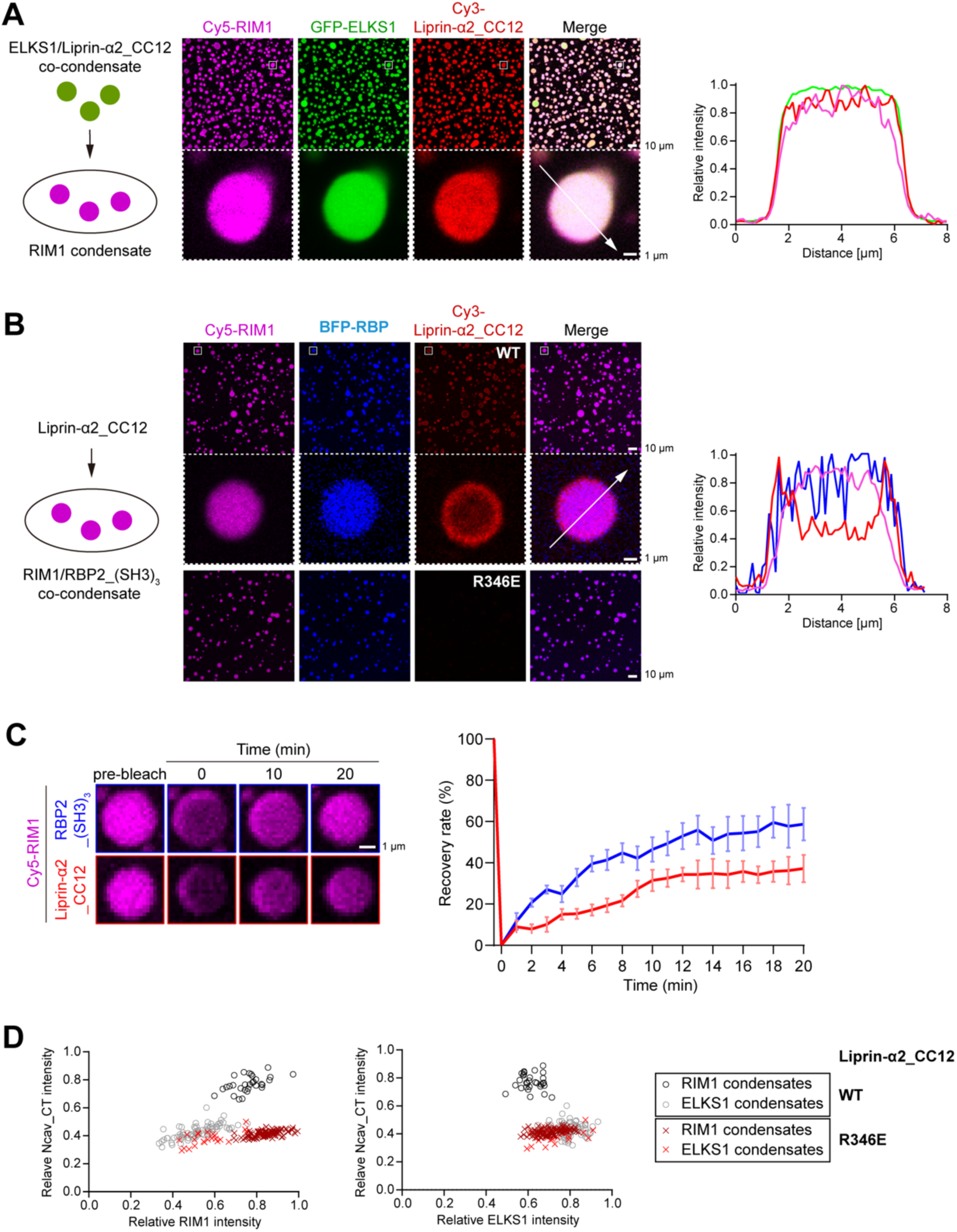
*In vitro* LLPS analyses of active zone proteins. **A.** Confocal imaging analysis of the LLPS mixture containing ELKS1/liprin-α_CC12 and RIM1 condensates. A magnified view of a representative droplet is displayed below, with a line analysis of fluorescence signal intensities along the indicated dashed line. The concentration of each protein was 5 μM. **B.** Confocal imaging analysis of RIM1/RBP2 co-condensates upon adding liprin-α_CC12 WT or the R346E mutant. A magnified view of a representative droplet, as boxed in the WT condition, is displayed below, with a line analysis of fluorescence signal intensities along the dashed line. The concentration of each protein was 5 μM. **C.** FRAP analysis of RIM1 condensates in the presence of RBP2_(SH3)_3_ or liprin-α_CC12. **D.** Plot analyses illustrating the intensity relationship between NCav_CT and RIM1 (left panel) or ELKS1 (right panel) fluorescence signals in RIM1 or ELKS1 condensates in the presence of liprin-α_CC12 WT or R346E.

**Table S1.** X-ray data collection and refinement statistics.

| Data collection |  |
| --- | --- |
| Space group | *P* 2<sub>1</sub> |
| Cell dimensions |  |
| *a*, *b*, *c* (Å) | 58.663, 92.697, 66.583 |
| α, β, γ (°) | 90, 102.95, 90 |
| Resolution (Å) | 50–2.75 (2.8–2.75) |
| *R*<sub>merge</sub><sup>a</sup> | 0.149 (1.151) |
| *I*/σ*I* | 13.8 (1.4) |
| *CC*<sub>1/2</sub><sup>b</sup> | 0.995 (0.714) |
| Completeness (%) | 98.9 (98.0) |
| Redundancy | 6.3 (5.9) |
| Refinement |  |
| Resolution (Å) | 50–2.75 (2.92–2.75) |
| No. reflections | 17881 (2950) |
| *R*<sub>work</sub> / *R*<sub>free</sub><sup>c</sup> | 0.205 (0.290) / 0.234 (0.323) |
| No. atoms |  |
| Protein | 3832 |
| Ligand/ion | 0 |
| Water | 20 |
| Mean *B* (Å) |  |
| Protein | 68.8 |
| Ligand/ion | - |
| Water | 58.9 |
| r.m.s. deviations |  |
| Bond lengths (Å) | 0.002 |
| Bond angles (°) | 0.5 |
| Ramachandran analysis |  |
| Favored region (%) | 98.7 |
| Allowed region (%) | 1.3 |
| Outliers (%) | 0 |

The numbers in parentheses represent values for the highest resolution shell.
<sup>a</sup>*R*<sub>merge</sub> = Σ|*I*<sub>i</sub> - *I*<sub>m</sub>|/Σ*I*<sub>i</sub>, where *I*<sub>i</sub> is the intensity of the measured reflection and *I*<sub>m</sub> is the mean intensity of all symmetry related reflections.
<sup>b</sup>*CC*<sub>1/2</sub> is the correlation coefficient of the half datasets.
<sup>c</sup>*R*<sub>work</sub> = Σ||*F*<sub>obs</sub>| - |*F*<sub>calc</sub>||/Σ|*F*<sub>obs</sub>|, where *F*<sub>obs</sub> and *F*<sub>calc</sub> are observed and calculated structure factors.
*R*<sub>free</sub> = Σ<sub>T</sub>||*F*<sub>obs</sub>| - |*F*<sub>calc</sub>||/Σ<sub>T</sub>|*F*<sub>obs</sub>|, where *T* is a test data set of about 5 % of the total reflections randomly chosen and set aside prior to refinement.

**Table S2.** Summary of plasmids.

| PLASMID | SOURCE / RRID |
| --- | --- |
| 32m3c-liprin-α2_CC2 (aa 259-542) | This paper |
| pET28a-RIM1α_C2B (aa 1166-1334) | This paper |
| pET28a-RIM1α_C2B (E1198R) | This paper |
| pET28a-RIM1α_C2B (R1201Q) | This paper |
| pET28a-RIM1α_C2B (R1239E) | This paper |
| pET28a-RIM1α_C2B (L1245Q) | This paper |
| pET28a-RIM1α_C2B (Q1247A) | This paper |
| pET28a-RIM1α_C2B (R1236E) | This paper |
| pET28a-RIM1α_C2B (K1240E) | This paper |
| 32m3c-liprin-α2_CC2C (aa 404-542) | This paper |
| 32m3c-liprin-α2_CC2N (aa 300-404) | This paper |
| 32m3c-liprin-α2_CC2N (E337R) | This paper |
| 32m3c-liprin-α2_CC2N (R339E) | This paper |
| 32m3c-liprin-α2_CC2N (E344R) | This paper |
| 32m3c-liprin-α2_CC2N (R346E) | This paper |
| 32m3c-liprin-α2_CC2N (L348F) | This paper |
| 32m3c-liprin-α2_CC2N (A350S) | This paper |
| 32m3c-liprin-α2_CC2N (E379R) | This paper |
| 32m3c-liprin-α2_CC2N (R383A) | This paper |
| 32m3c-liprin-α2_CC2N (E380R) | This paper |
| pET28a-liprin-α2_CC12 (aa 24-542) | Liang et al., 2021 |
| pET28a-liprin-α2_CC12 (E344R) | This paper |
| pET28a-liprin-α2_CC12 (R346E) | This paper |
| pET28a-liprin-α2_CC12 (R383A) | This paper |
| m3c-RBP2_(SH3)<sub>3</sub> (aa 178-252+844-1040) | Wu et al., 2019 |
| 32m3c-NCav_CT (aa 2151-2327) | Wu et al., 2019 |
| pCAG-FLAG-RIM1 | This paper |
| pCAG-FLAG-ELKS1 | This paper |
| pTGFP-Liprin-α2 | Xie et al., 2020; |
| pTGFP-Liprin-α2-ΔCC2N (aa 1-299+405-1257) | This paper |
| pTGFP-Liprin-α2-ΔCC2C (aa 1-404+543-1257) | This paper |
| pTGFP-Liprin-α2 (E337R) | This paper |
| pTGFP-Liprin-α2 (R339E) | This paper |
| pTGFP-Liprin-α2 (E344R) | This paper |
| pTGFP-Liprin-α2 (R346E) | This paper |
| pFU-EGFP-Liprin-α2 (WT) | This paper |
| pFU-EGFP-Liprin-α2 (ΔCC2N) | This paper |
| pFU-EGFP-Liprin-α2 (E344R) | This paper |
| pFU-EGFP-Liprin-α2 (R346E) | This paper |
| pFU-NLS-GFP-Cre | Marcó de la Cruz et al., 2024 |
| pMDLg/pRRE | RRID: Addgene_12251 |
| pRSV-REV | RRID: Addgene_12253 |
| pVSVG | RRID: Addgene_35616 |
| pFU-M2rtTA | RRID: Addgene_20342 |
| pTet-O-Ngn2-puromycin | RRID: Addgene_52047 |
| pFU-oChIEF-tdTomato | Marcó de la Cruz et al., 2024 |

**Table S3.**
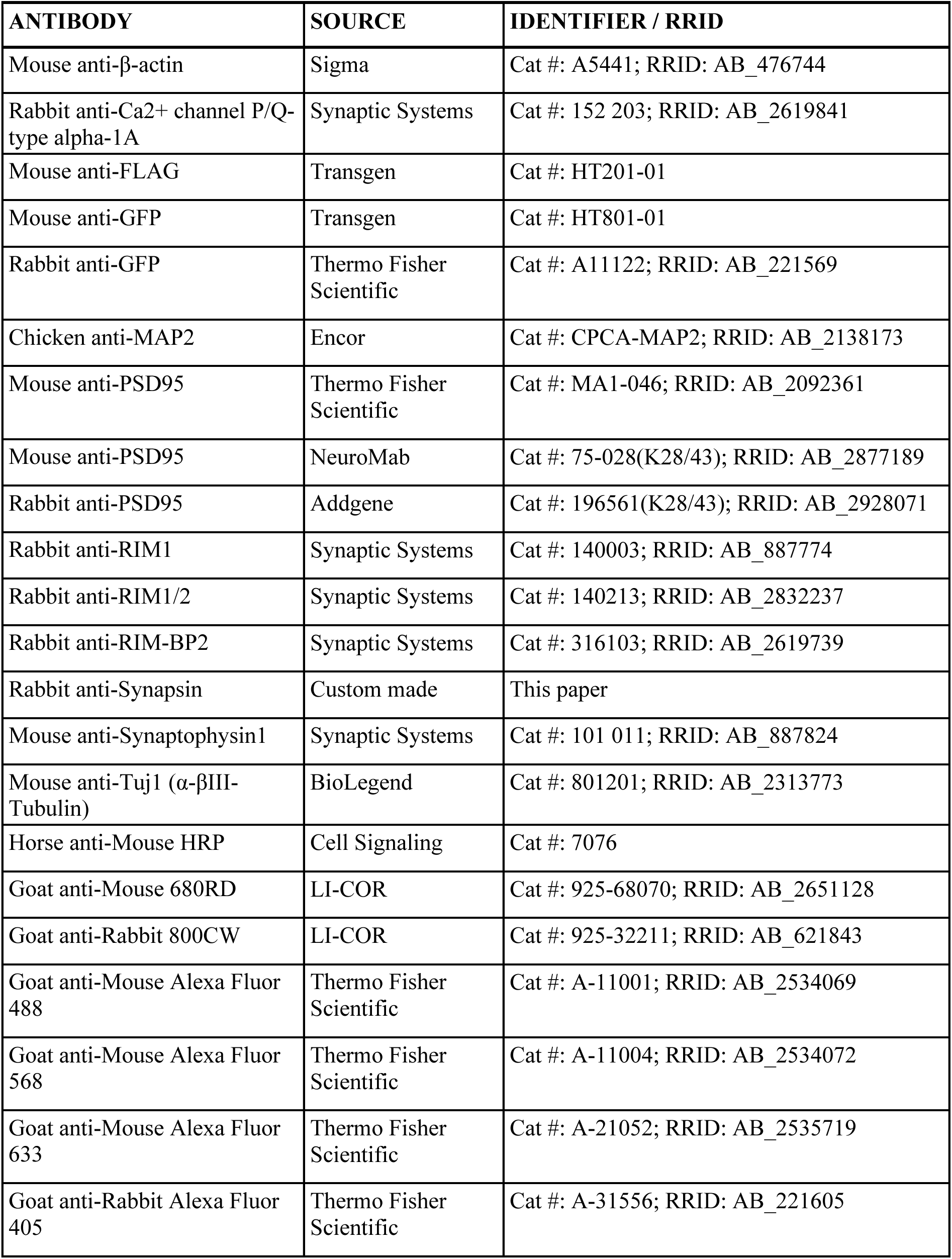
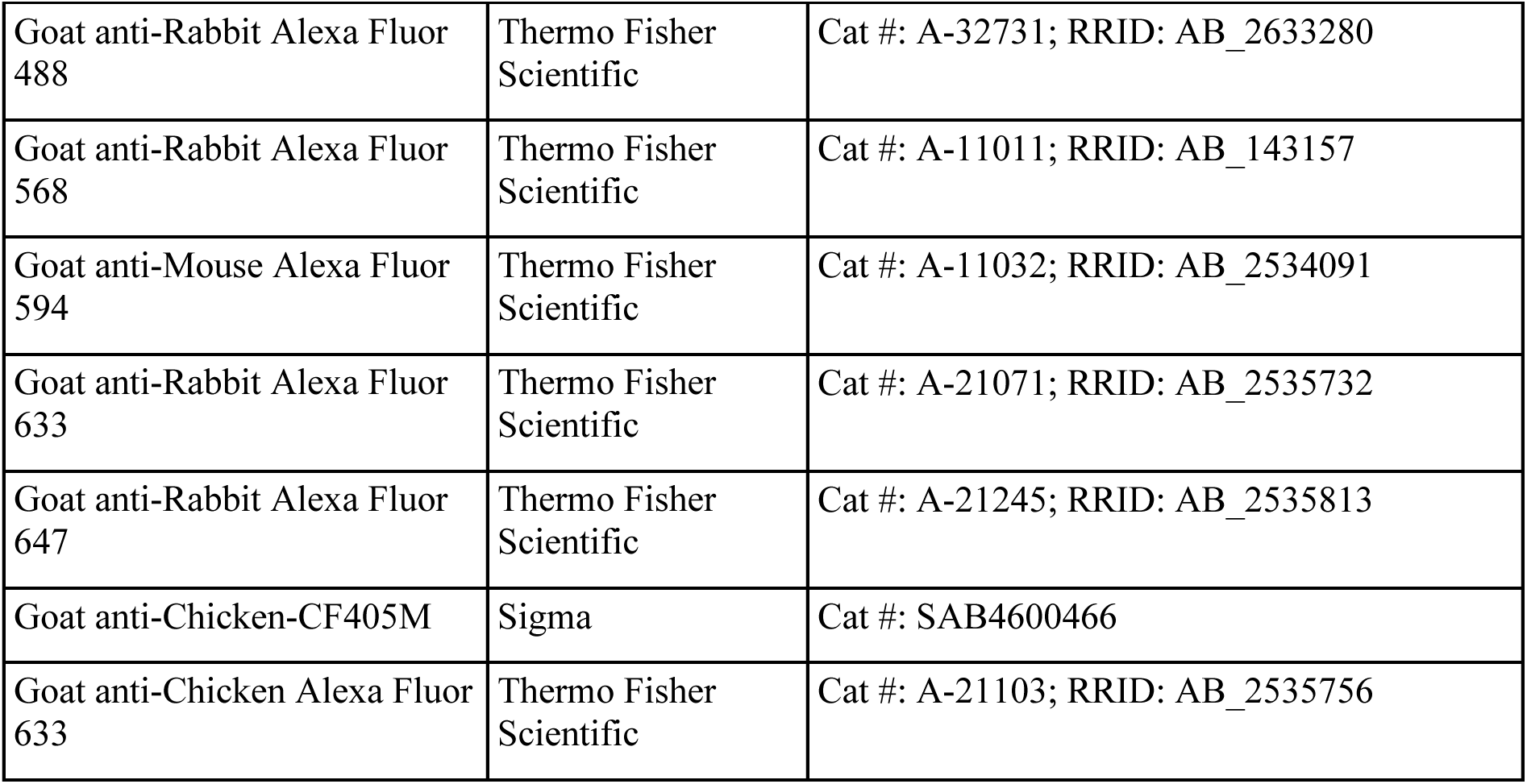
Antibody identifiers.

| ANTIBODY | SOURCE | IDENTIFIER / RRID |
| --- | --- | --- |
| Mouse anti-β-actin | Sigma | Cat #: A5441; RRID: AB_476744 |
| Rabbit anti-Ca<sup>2+</sup> channel P/Q-type α-1A | Synaptic Systems | Cat #: 152 203; RRID: AB_2619841 |
| Mouse anti-FLAG | Transgen | Cat #: HT201-01 |
| Mouse anti-GFP | Transgen | Cat #: HT801-01 |
| Rabbit anti-GFP | Thermo Fisher Scientific | Cat #: A11122; RRID: AB_221569 |
| Chicken anti-MAP2 | Encor | Cat #: CPCA-MAP2; RRID: AB_2138173 |
| Mouse anti-PSD95 | Thermo Fisher Scientific | Cat #: MA1-046; RRID: AB_2092361 |
| Mouse anti-PSD95 | NeuroMab | Cat #: 75-028(K28/43); RRID: AB_2877189 |
| Rabbit anti-PSD95 | Addgene | Cat #: 196561(K28/43); RRID: AB_2928071 |
| Rabbit anti-RIM1 | Synaptic Systems | Cat #: 140003; RRID: AB_887774 |
| Rabbit anti-RIM1/2 | Synaptic Systems | Cat #: 140213; RRID: AB_2832237 |
| Rabbit anti-RIM-BP2 | Synaptic Systems | Cat #: 316103; RRID: AB_2619739 |
| Rabbit anti-Synapsin | Custom made | This paper |
| Mouse anti-Synaptophysin1 | Synaptic Systems | Cat #: 101 011; RRID: AB_887824 |
| Mouse anti-Tuj1 (α-βIII-Tubulin) | BioLegend | Cat #: 801201; RRID: AB_2313773 |
| Horse anti-Mouse HRP | Cell Signaling | Cat #: 7076 |
| Goat anti-Mouse 680RD | LI-COR | Cat #: 925-68070; RRID: AB_2651128 |
| Goat anti-Rabbit 800CW | LI-COR | Cat #: 925-32211; RRID: AB_621843 |
| Goat anti-Mouse Alexa Fluor 488 | Thermo Fisher Scientific | Cat #: A-11001; RRID: AB_2534069 |
| Goat anti-Mouse Alexa Fluor 568 | Thermo Fisher Scientific | Cat #: A-11004; RRID: AB_2534072 |
| Goat anti-Mouse Alexa Fluor 633 | Thermo Fisher Scientific | Cat #: A-21052; RRID: AB_2535719 |
| Goat anti-Rabbit Alexa Fluor 405 | Thermo Fisher Scientific | Cat #: A-31556; RRID: AB_221605 |
| Goat anti-Rabbit Alexa Fluor 488 | Thermo Fisher Scientific | Cat #: A-32731; RRID: AB_2633280 |
| Goat anti-Rabbit Alexa Fluor 568 | Thermo Fisher Scientific | Cat #: A-11011; RRID: AB_143157 |
| Goat anti-Mouse Alexa Fluor 594 | Thermo Fisher Scientific | Cat #: A-11032; RRID: AB_2534091 |
| Goat anti-Rabbit Alexa Fluor 633 | Thermo Fisher Scientific | Cat #: A-21071; RRID: AB_2535732 |
| Goat anti-Rabbit Alexa Fluor 647 | Thermo Fisher Scientific | Cat #: A-21245; RRID: AB_2535813 |
| Goat anti-Chicken-CF405M | Sigma | Cat #: SAB4600466 |
| Goat anti-Chicken Alexa Fluor 633 | Thermo Fisher Scientific | Cat #: A-21103; RRID: AB_2535756 |

## Notes

### Competing Interest Statement

The authors have declared no competing interest.

## References

1. Südhof, T. C. The presynaptic active zone. Neuron 75, 11–25 (2012).

2. Ackermann, F., Waites, C. L. & Garner, C. C. Presynaptic active zones in invertebrates and vertebrates. EMBO Rep 16, 1–16 (2015).

3. Emperador-Melero, J. & Kaeser, P. S. Assembly of the presynaptic active zone. Curr Opin Neurobiol 63, 95–103 (2020).

4. Serra-Pages, C., Medley, Q. G., Tang, M., Hart, A. & Streuli, M. Liprins, a family of LAR transmembrane protein-tyrosine phosphatase-interacting proteins. J Biol Chem 273, 15611– 15620 (1998).

5. Zhen, M. & Jin, Y. The liprin protein SYD-2 regulates the differentiation of presynaptic termini in C. elegans. Nature 401, 371–375 (1999).

6. Kaufmann, N., DeProto, J., Ranjan, R., Wan, H. & Van Vactor, D. Drosophila liprin-alpha and the receptor phosphatase Dlar control synapse morphogenesis. Neuron 34, 27–38 (2002).

7. Spangler, S. A. et al. Differential expression of liprin-alpha family proteins in the brain suggests functional diversification. J Comp Neurol 519, 3040–3060 (2011).

8. Zürner, M. et al. Analyses of the spatiotemporal expression and subcellular localization of liprin-alpha proteins. J Comp Neurol 519, 3019–3039 (2011).

9. Kittelmann, M. et al. Liprin-α/SYD-2 determines the size of dense projections in presynaptic active zones in C. elegans. J Cell Biol 203, 849–63 (2013).

10. Wong, M. Y. et al. Liprin-α3 controls vesicle docking and exocytosis at the active zone of hippocampal synapses. Proceedings of the National Academy of Sciences 115, 2234–2239 (2018).

11. Marcó de la Cruz, B., et al. Liprin-α proteins are master regulators of human presynapse assembly. Nat Neurosci 27, 629–642 (2024).

12. Ackley, B. D. et al. The two isoforms of the Caenorhabditis elegans leukocyte-common antigen related receptor tyrosine phosphatase PTP-3 function independently in axon guidance and synapse formation. J Neurosci 25, 7517–7528 (2005).

13. Olsen, O. et al. Neurotransmitter release regulated by a MALS-liprin-alpha presynaptic complex. J Cell Biol 170, 1127–1134 (2005).

14. Patel, M. R. et al. Hierarchical assembly of presynaptic components in defined C. elegans synapses. Nat Neurosci 9, 1488–1498 (2006).

15. Spangler, S. A. et al. Liprin-alpha2 promotes the presynaptic recruitment and turnover of RIM1/CASK to facilitate synaptic transmission. J Cell Biol 201, 915–928 (2013).

16. Xie, X., Liang, M., Yu, C. & Wei, Z. Liprin-α-Mediated Assemblies and Their Roles in Synapse Formation. Front Cell Dev Biol 9, (2021).

17. Xie, X. et al. Structural basis of liprin-α-promoted LAR-RPTP clustering for modulation of phosphatase activity. Nat Commun 11, 169 (2020).

18. Wei, Z. et al. Liprin-mediated large signaling complex organization revealed by the liprin-alpha/CASK and liprin-alpha/liprin-beta complex structures. Mol Cell 43, 586–598 (2011).

19. Wakita, M. et al. Structural insights into selective interaction between type IIa receptor protein tyrosine phosphatases and Liprin-α. Nat Commun 11, 649 (2020).

20. Schoch, S. et al. RIM1alpha forms a protein scaffold for regulating neurotransmitter release at the active zone. Nature 415, 321–326 (2002).

21. Ko, J., Na, M., Kim, S., Lee, J.-R. R. & Kim, E. Interaction of the ERC family of RIM-binding proteins with the liprin-alpha family of multidomain proteins. J Biol Chem 278, 42377–42385 (2003).

22. Dai, Y. et al. SYD-2 Liprin-alpha organizes presynaptic active zone formation through ELKS. Nat Neurosci 9, 1479–1487 (2006).

23. Liang, M. et al. Oligomerized liprin-α promotes phase separation of ELKS for compartmentalization of presynaptic active zone proteins. Cell Rep 34, 108901 (2021).

24. Ohtsuka, T. et al. Cast: a novel protein of the cytomatrix at the active zone of synapses that forms a ternary complex with RIM1 and munc13-1. J Cell Biol 158, 577–590 (2002).

25. Wang, Y., Liu, X., Biederer, T. & Südhof, T. C. A family of RIM-binding proteins regulated by alternative splicing: Implications for the genesis of synaptic active zones. Proceedings of the National Academy of Sciences 99, 14464–14469 (2002).

26. Kaeser, P. S. et al. RIM Proteins Tether Ca2+ Channels to Presynaptic Active Zones via a Direct PDZ-Domain Interaction. Cell 144, 282–295 (2011).

27. Lu, J. et al. Structural basis for a Munc13-1 homodimer to Munc13-1/RIM heterodimer switch. PLoS Biol 4, 1159–1172 (2006).

28. Wang, Y. & Südhof, T. C. Genomic definition of RIM proteins: Evolutionary amplification of a family of synaptic regulatory proteins. Genomics 81, (2003).

29. de Jong, A. P. H. et al. RIM C2B Domains Target Presynaptic Active Zone Functions to PIP2-Containing Membranes. Neuron 98, 335–349.e7 (2018).

30. Wu, X. et al. RIM and RIM-BP Form Presynaptic Active-Zone-like Condensates via Phase Separation. Mol Cell 73, 971–984.e5 (2019).

31. Wu, X. et al. Vesicle Tethering on the Surface of Phase-Separated Active Zone Condensates. Mol Cell 81, 13–24.e7 (2021).

32. McDonald, N. A., Fetter, R. D. & Shen, K. Assembly of synaptic active zones requires phase separation of scaffold molecules. Nature 588, 454–458 (2020).

33. Paul, M. S. et al. A syndromic neurodevelopmental disorder caused by rare variants in PPFIA3. Am J Hum Genet 111, 96–118 (2024).

34. Sourbron, J., Jansen, K., Aerts, N. & Lagae, L. PPFIA4 mutation: A second hit in POLG related disease? Epilepsy Behav Rep 16, 100455 (2021).

35. Iossifov, I. et al. The contribution of de novo coding mutations to autism spectrum disorder. Nature 515, 216–221 (2014).

36. Reuter, M. S. et al. Diagnostic yield and novel candidate genes by exome sequencing in 152 consanguineous families with neurodevelopmental disorders. JAMA Psychiatry 74, 293– 299 (2017).

37. Kaplanis, J. et al. Evidence for 28 genetic disorders discovered by combining healthcare and research data. Nature 586, (2020).

38. Guan, R. et al. Crystal structure of the RIM1α C2B domain at 1.7 Å resolution. Biochemistry 46, 8988–8998 (2007).

39. Emperador-Melero, J. et al. PKC-phosphorylation of Liprin-α3 triggers phase separation and controls presynaptic active zone structure. Nat Commun 12, (2021).

40. Banani, S. F., Lee, H. O., Hyman, A. A. & Rosen, M. K. Biomolecular condensates: organizers of cellular biochemistry. Nat Rev Mol Cell Biol 18, 285–298 (2017).

41. Chen, X., Wu, X., Wu, H. & Zhang, M. Phase separation at the synapse. Nat Neurosci 23, 301–310 (2020).

42. Rosenmund, C. & Stevens, C. F. Definition of the readily releasable pool of vesicles at hippocampal synapses. Neuron 16, (1996).

43. Betz, A. et al. Functional interaction of the active zone proteins Munc13-1 and RIM1 in synaptic vesicle priming. Neuron 30, 183–196 (2001).

44. Koushika, S. P. et al. A post-docking role for active zone protein Rim. Nat Neurosci 4, 997– 1005 (2001).

45. Han, Y., Kaeser, P. S., Südhof, T. C. & Schneggenburger, R. RIM determines Ca2+ channel density and vesicle docking at the presynaptic active zone. Neuron 69, 304–316 (2011).

46. Acuna, C., Liu, X., Gonzalez, A. & Südhof, T. C. RIM-BPs Mediate Tight Coupling of Action Potentials to Ca2+-Triggered Neurotransmitter Release. Neuron 87, (2015).

47. Dong, W. et al. CAST/ELKS Proteins Control Voltage-Gated Ca2+ Channel Density and Synaptic Release Probability at a Mammalian Central Synapse. Cell Rep 24, 284–293.e6 (2018).

48. Lu, J., Li, H., Wang, Y., Südhof, T. C. & Rizo, J. Solution structure of the RIM1α PDZ domain in complex with an ELKS1b C-terminal peptide. J Mol Biol 352, 455–466 (2005).

49. Tang, A.-H. et al. A trans-synaptic nanocolumn aligns neurotransmitter release to receptors. Nature 536, 210–214 (2016).

50. Trotter, J. H. et al. Synaptic neurexin-1 assembles into dynamically regulated active zone nanoclusters. J Cell Biol 218, 2677–2698 (2019).

51. Sakamoto, H. et al. Synaptic weight set by Munc13-1 supramolecular assemblies. Nat Neurosci 21, 41–49 (2018).

52. Ghelani, T. et al. Interactive nanocluster compaction of the ELKS scaffold and Cacophony Ca2+ channels drives sustained active zone potentiation. Sci Adv 9, (2023).

53. Rebola, N. et al. Distinct Nanoscale Calcium Channel and Synaptic Vesicle Topographies Contribute to the Diversity of Synaptic Function. Neuron 104, 693–710.e9 (2019).

54. Patel, M. R. & Shen, K. RSY-1 is a local inhibitor of presynaptic assembly in C. elegans. Science (1979) 323, 1500–1503 (2009).

55. Ferrer-Orta, C. et al. Structural characterization of the Rabphilin-3A-SNAP25 interaction. Proc Natl Acad Sci U S A 114, E5343–E5351 (2017).

56. Zhou, Q. et al. The primed SNARE–complexin–synaptotagmin complex for neuronal exocytosis. Nature 548, 420–425 (2017).

57. Guillén, J. et al. Structural insights into the Ca2+ and PI(4,5)P2 binding modes of the C2 domains of rabphilin 3A and synaptotagmin 1. Proc Natl Acad Sci U S A 110, 20503–20508 (2013).

58. Wang, J. et al. Circular oligomerization is an intrinsic property of synaptotagmin. Elife 6, (2017).

59. Honigmann, A. et al. Phosphatidylinositol 4,5-bisphosphate clusters act as molecular beacons for vesicle recruitment. Nat Struct Mol Biol 20, 679–686 (2013).

60. Wang, S., Li, Y. & Ma, C. Synaptotagmin-1 C2B domain interacts simultaneously with SNAREs and membranes to promote membrane fusion. Elife 5, (2016).

61. Jin, G. et al. Structural basis of ELKS/Rab6B interaction and its role in vesicle capturing enhanced by liquid-liquid phase separation. Journal of Biological Chemistry 299, 104808 (2023).

62. Otwinowski, Z. & Minor, W. Processing of X-ray Diffraction Data Collected in Oscillation Mode. Methods Enzymol 276, 307–326 (1997).

63. Storoni, L. C., McCoy, A. J. & Read, R. J. Likelihood-enhanced fast rotation functions. Acta Crystallogr D Biol Crystallogr 60, 432–438 (2004).

64. Emsley, P. & Cowtan, K. Coot: model-building tools for molecular graphics. Acta Crystallogr D Biol Crystallogr 60, 2126–2132 (2004).

65. Adams, P. D. et al. PHENIX: A comprehensive Python-based system for macromolecular structure solution. Acta Crystallogr D Biol Crystallogr 66, 213–221 (2010).

66. Davis, I. W. et al. MolProbity: all-atom contacts and structure validation for proteins and nucleic acids. Nucleic Acids Res 35, W375–83 (2007).

67. Sterky, F. H. et al. Carbonic anhydrase-related protein CA10 is an evolutionarily conserved pan-neurexin ligand. Proc Natl Acad Sci U S A 114, (2017).

68. Zhang, Y. et al. Rapid single-step induction of functional neurons from human pluripotent stem cells. Neuron 78, (2013).

69. Tan, C. et al. Munc13 supports fusogenicity of non-docked vesicles at synapses with disrupted active zones. Elife 11, (2022).

